# Reduction of *Grin2a* in adolescent rat dopamine neurons confers a phenotype relevant to psychosis

**DOI:** 10.1101/2024.10.28.620713

**Authors:** Michelle L. Kielhold, David S. Jacobs, Alejandro Torrado-Pacheco, Merridee J. Lefner, Joseph J. Lebowitz, Angela J. Langdon, John T. Williams, Larry S. Zweifel, Bita Moghaddam

**Author notes:** Corresponding author: Bita Moghaddam.

## Abstract

Psychosis is a hallmark of schizophrenia. It involves a collection of symptoms that are typically associated with disrupted dopamine signaling and emerges during adolescence or early adulthood. Most schizophrenia-associated genes, however, involve glutamatergic or other ubiquitous targets that do not explain the latent expression of psychosis or dopaminergic abnormalities. Here, we describe an etiologically relevant model for the adolescent onset of dopamine-related dysfunction in schizophrenia. We focused on *GRIN2A,* the gene encoding the GluN2A subunit of the NMDA receptor, as both the common loss-of-function variants and the rare missense variants in this gene are risk factors for schizophrenia. We find that GluN2A levels distinctly decline in dopamine neuron-containing regions throughout adolescence while remaining stable in other regions. This suggested that adolescent dopamine neurons may be particularly vulnerable to further reductions in GluN2A caused by a damaging variant of *GRIN2A*. Consistent with this idea, we find that selective knockout of *Grin2a* in adolescent rat dopamine neurons results in a psychosis-relevant behavioral phenotype. This manipulation also reduced dopamine release in response to unexpected outcomes in young adults, reflective of prediction error signaling abnormalities observed in the clinical population. These data provide mechanistic insight into how *GRIN2A* mutations may contribute to the delayed onset of dopamine-related symptoms and provide a model for identifying course altering treatments for schizophrenia.

## INTRODUCTION

NMDA receptor-mediated glutamatergic neurotransmission has long been implicated in the pathophysiology of schizophrenia (SCZ)^1,2^. This association initially stemmed from postmortem findings^3–5^ and the discovery that the drug phencyclidine, which transiently mimics SCZ symptoms, is an antagonist of the NMDA receptor^6,7^. Recent genetic findings have strengthened evidence implicating the NMDA receptor in the etiology of SCZ and, more specifically, have directed the focus toward the GluN2A subunit of this receptor. Both common and rare variants that reduce the function of the gene encoding this subunit (*GRIN2A*) have been identified as genetic risk factors for SCZ^8–13^. In line with this discovery, *Grin2a* mutant mice display a wide range of aberrant cortical activity and locomotor abnormalities^14–16^.

Dopamine neurotransmission is equally implicated in SCZ pathophysiology and psychosis^17–19^. Notably, all known antipsychotic drugs act on dopamine receptors^20–23^. While genetic vulnerability to develop SCZ generally does not suggest a primary role for dopamine receptors or related proteins, extensive imaging and pharmacological studies have implicated the dopamine system in the positive symptoms of the disorder. PET studies, for example, have reported disrupted dopamine neurotransmission in individuals with SCZ or young, at-risk individuals who go on to develop the disorder^24–26^.

The involvement of *GRIN2A* in the etiology of SCZ may explain its distinct temporal pattern of symptom expression. SCZ is characterized by three types of symptoms: cognitive, affective (negative), and psychotic (positive) symptoms. Cognitive and affective symptoms often emerge in late childhood, whereas psychotic symptoms typically manifest in late adolescence or early adulthood and are associated with disruptions in dopamine neurotransmission^17^. In humans and rodents *GRIN2A/Grin2a* cortical expression is absent or undetected early in development but steeply increases with age, beginning perinatally (humans) or after birth (rodents), reaching adult levels in early childhood^27–31^. Thus, phenotypes related to a reduction in functional *GRIN2A* would emerge during this early developmental stage, which coincides with the timeline of cognitive and affective symptom expression in SCZ. But why would psychosis remain latent until late adolescence or early adulthood? Moreover, why are these late-emerging positive symptoms—and not the negative or cognitive symptoms—alleviated by dopamine receptor antagonists^21–25,32–38^. These questions led us to investigate how reduced function of *GRIN2A* in adolescent dopamine neurons may lead to presentation of behavioral phenotypes that are relevant to psychosis.

We found that, unlike cortical regions where GluN2A remains stable during adolescence, levels in dopamine neuron-containing regions decline at this age. This suggested that a loss of function in *Grin2a* may exaggerate this natural decline and be most detrimental to dopamine neurotransmission during adolescence. Consistent with this mechanism, targeted knockout of *Grin2a* in adolescent dopamine neurons was sufficient to produce a selective phenotype relevant to aspects of psychosis, without affecting spontaneous or simple reward seeking behaviors.

## RESULTS

### GluN2A expression is dynamic across postnatal development in a region-specific manner

Brain regions including the prefrontal cortex and hippocampus do not express GluN2A in rodents at birth, but levels rapidly increase postnatally, reaching adult levels around weaning^30,31,39^. It is, however, not known if GluN2A levels remain stable between weaning and adulthood. We quantified GluN2A protein levels in male and female rats immediately after weaning on postnatal day 21 (P21), during adolescence (P35, P45), and adulthood (P80) (**Figure 1a**), focusing on the medial prefrontal cortex (mPFC), hippocampus, and regions containing dopamine neurons (i.e. ventral tegmental area (VTA) and substantia nigra (SN)). We found that GluN2A levels in the mPFC and hippocampus remain stable between ages P21 and P80 (**Figure 1b**). In contrast, GluN2A levels were labile in dopamine neuron-rich regions during adolescence. In both the VTA and SN, GluN2A levels declined after P21, reaching adult levels during mid-to-late adolescence (**Figure 1c**).

**Figure 1:**
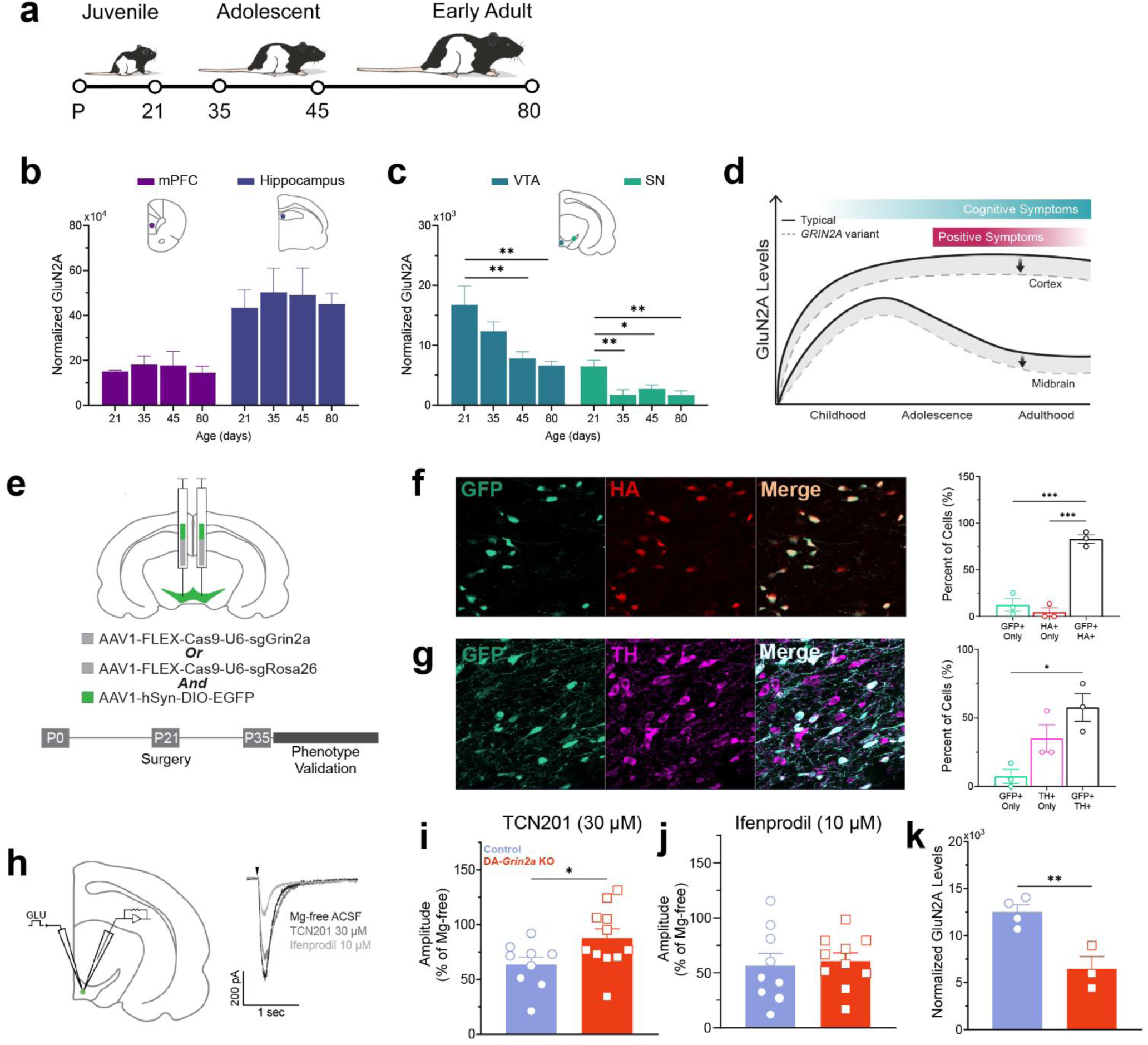
GluN2A loss in the adolescent midbrain was augmented with virally mediated *Grin2a* selective knockout in dopamine neurons. **a**, Timeline of tissue collection ages for postnatal day 21 (P21), adolescent (P35, P45), and adult (P80) samples. **b,** Medial prefrontal cortex (mPFC) and hippocampus showed no effect of age on normalized GluN2A levels. **c,** Ventral tegmental area (VTA) showed significant effect of age, with reductions in normalized GluN2A at P21 vs P45 (\**p*=0.0157) and vs P80 (\*\**p*=0.0058). The substantia nigra (SN) also showed an effect of age with reductions from P21 vs P35 (\*\**p*=0.0036), vs P45 (\**p*=0.0318), and vs P80 (\*\**p*=0.0033); (*n*=5-6 brains). Significance testing: one-way ANOVA; Dunnett post hoc. **d,** Representation of correlation between adolescent emergence of psychosis and midbrain GluN2A loss. **e,** Schematic representation of virus infusion protocol and timeline of phenotyping. **f,** Representative image of GFP overlap with hemagglutinin (HA)-tagged CRISPR/Cas9 virus, quantification shows significantly more cells expressed both GFP and HA than only GFP (\*\*\**p*=0.0002) or only HA (\*\*\**p*=0.0001) **g,** Representative image showing GFP overlap with dopaminergic cells, quantification shows significantly more cells expressed both GFP and tyrosine hydroxylase (TH) than only GFP (\**p*=0.0152); (*n*=3 brains). Significance testing: one-way ANOVA; Tukey post hoc. **h,** Schematic representation of experimental design and example traces of slice electrophysiology experiments in adolescent brain. **i,** Quantification of peak amplitudes showed dopamine cells in DA-*Grin2a* KO brains were significantly less affected by TCN201 than controls (\**p*=0.049); (control *n*=9, DA-*Grin2a* KO *n*=11 cells). **j,** Quantification of peak amplitudes showed dopamine cells in DA-*Grin2a* KO brains were similarly affected by ifenprodil as controls; (control *n*=9, DA-*Grin2a* KO *n*=10 cells). **k**, Western blot analysis of adult brain tissue showed a significant reduction of normalized GluN2A in the VTA of DA-*Grin2a* KO animals compared to controls (\*\**p* =0.0082); (control *n*=4, DA-*Grin2a* KO *n*=3 brains). Significance testing: unpaired t-test. Error bars denote ±SEM.

This observation led us to hypothesize that the damaging variants of *GRIN2A* implicated in SCZ may exacerbate this natural reduction of GluN2A in dopamine neurons during adolescence, providing a mechanism for the emergence of psychosis at this developmental stage (**Figure 1d**). We tested this idea by establishing a model of adolescent dopamine neuron-selective knockout of *Grin2a* (DA-*Grin2a* KO) through the infusion of a Cre-inducible CRISPR/Cas9 virus targeting *Grin2a* into the VTA of TH::Cre rats. This virus was co-infused with a Cre-dependent virus that would express green fluorescent protein (GFP) for visualization (**Figure 1e**). Viral infusions were performed on the day of weaning (P21) and expression was assessed following a 2-week incubation period.

Immunohistochemistry confirmed cell-specific virus expression, as staining for the hemagglutinin (HA)-tag on the DA-*Grin2a* KO virus significantly overlapped with GFP (**Figure 1f**) and this expression was specific to dopamine neurons as there was significant co-expression of GFP and tyrosine hydroxylase (TH) (**Figure 1g**). Functional reduction of GluN2A in GFP+ neurons was established with recordings in slice preparation (**Figure 1h**).

Bath application of a GluN2A-specific antagonist (TCN201) resulted in a reduction of the glutamate-induced current in controls whereas DA-*Grin2a* KO cells showed a significantly blunted response (**Figure 1i**). This reduction was specific to GluN2A-containing NMDA receptors as a GluN2B-targeting antagonist (ifenprodil) produced similar effects in cells of both control and DA-*Grin2a* KO animals (**Figure 1j**). The extent of this regional loss of GluN2A was determined through Western blot analysis, where we observed an approximate 50% reduction of GluN2A in the VTA of DA-*Grin2a* KO animals compared to controls (**Figure 1k**).

### DA-Grin2a KO adolescents retain typical spontaneous behaviors but have hypersensitivity to amphetamine

To assess the effect of DA-*Grin2a* KO on spontaneous unrewarded behaviors, a battery of tests was run to quantify spatial working memory, innate anxiety, and locomotion in adolescent animals (P45-55). Spontaneous alternation in a short-walled plus maze was used to assess spatial working memory. Since rats tend to explore novel arms over previously visited ones, impairments in working memory would emerge as reduced alternation in this task^40^. There were no significant between-group differences in spontaneous alternation in the plus maze, suggesting no effect of the DA-*Grin2a* KO on this form of spatial working memory in adolescents (**Figure 2a**). This was not due to differences in overall exploration of the maze as total arm entries were similar between groups (**Figure 2b**). In the elevated plus maze (EPM), avoidance of the open arms of the maze and preference for the walled-in, closed arms can be used to ascertain anxiety-like behavior in rodents^41^. There were no differences in anxiety-like behavior in the EPM as DA-*Grin2a* KO adolescents spent a similar percentage of time in the open and closed arms compared to controls (**Figure 2c,d**). Similar to EPM, avoidance of the exposed center portion of the open field maze is related to anxiety in rodents^42^. With this outcome measure, control and DA-*Grin2a* KO rats showed no significant difference in anxiety-like behavior in the open field task (**Figure 2e**). This was not a result of an impaired locomotor ability, as both treatment groups traveled similar distances in the open field (**Figure 2f**). We then performed a pharmacological challenge with amphetamine to determine the effect of the psychostimulant on locomotion. After the baseline period in the open field, animals received an injection of 1.0 mg/kg amphetamine and were returned to the field to determine their responsivity to the locomotor effects of the drug (**Figure 2g**). Amphetamine elicits hyperlocomotion by increasing dopamine levels, and hyperreactivity to the locomotor effects of this drug is considered a behavioral correlate of psychosis^43,44^. When we analyzed distance traveled in the open field under amphetamine in 10-min bins, we found DA-*Grin2a* KO rats displayed hypersensitivity to amphetamine (**Figure 2h**). Thus while “baseline” spontaneous behavior was not affected in DA-*Grin2a* KO animals, a difference emerged when the system was challenged by amphetamine.

**Figure 2:**
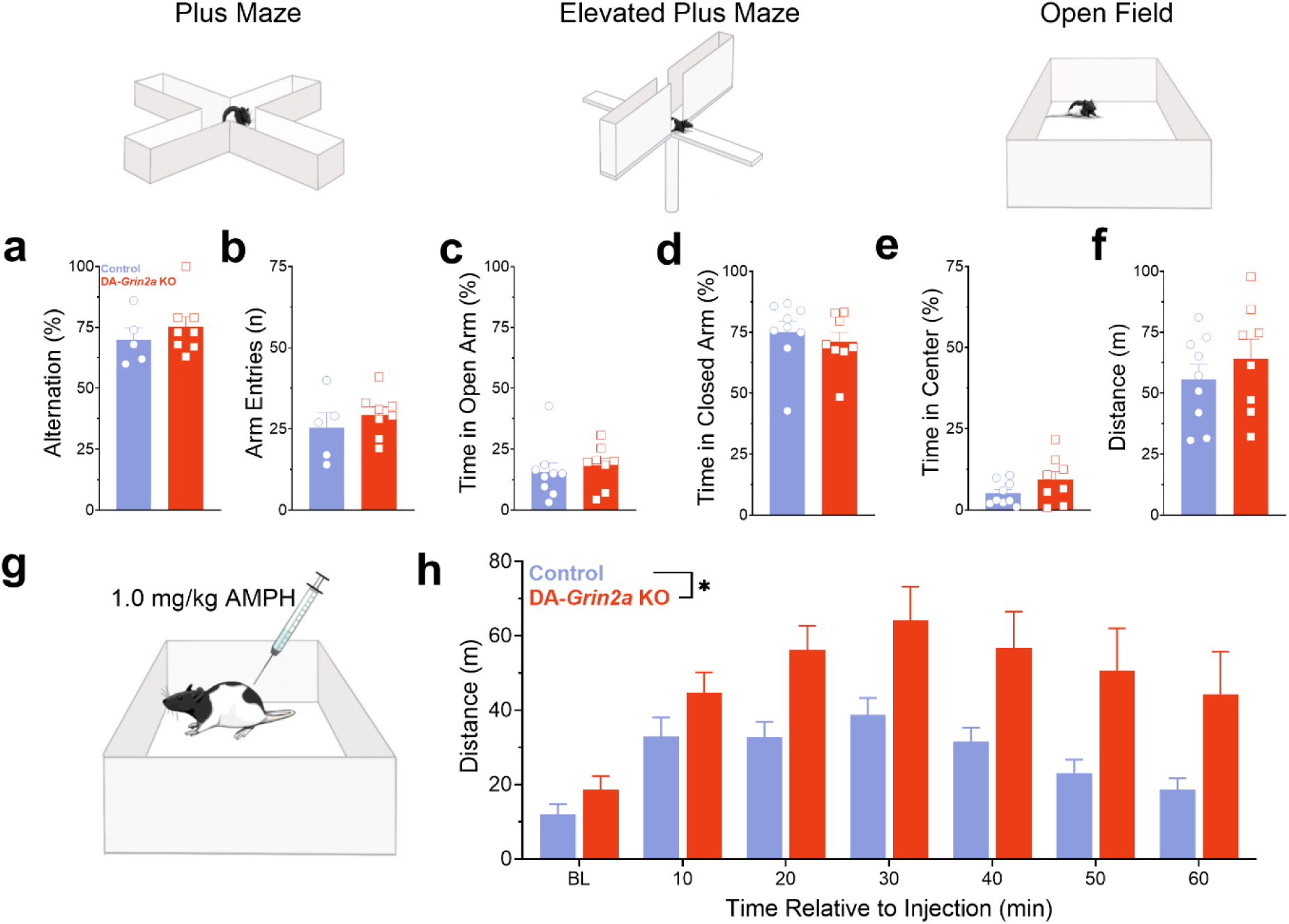
DA-*Grin2a* KO adolescents maintain typical spontaneous maze behaviors but show hypersensitivity to amphetamine. **a, b**, DA-*Grin2a* KO animals had no significant differences from controls in percent alternation or total arm entries in the plus maze; (control *n*=5, DA-*Grin2a* KO *n*=8 rats). **c, d**, Control and DA-*Grin2a* KO animals showed similar percent of the time spent in either the open or closed arms of the Elevated Plus Maze; (control *n*=9, DA-*Grin2a* KO *n*=8 rats). **e, f,** DA-*Grin2a* KO animals showed no significant differences from control animals in time spent in the center of the open field or in general locomotor activity in this maze; (control *n*=9, DA-*Grin2a* KO *n*=8 rats). Significance testing: unpaired t-test. **g,** Schematic depicting amphetamine (AMPH) administration in the open field. **h,** Distance traveled in 10 min time bins showed overall significant effect of treatment (*p=0.0397) with DA-*Grin2a* KO animals having higher distance traveled than controls; (control *n*=11, DA-*Grin2a* KO *n*=17 rats). Baseline (BL) bin was determined from the final 10 min of open field testing in (f). Significant testing: two-way ANOVA. Error bars denote ± SEM.

### DA-Grin2a KO disrupts adolescent reward-guided behavior during operant and Pavlovian conditioning in a construct-selective manner

The dopamine system is implicated in effort-, reward-, and punishment-related learning and associated behaviors. We used two distinct conditioning paradigms to assess the effect of *DA-Grin2a* KO on these learned behaviors.

#### Effort related operant behavior

Abnormalities in effort expenditure during reward seeking are a clinically observed phenomena in people with SCZ. When asked to perform high or low effort tasks for reward, individuals with SCZ show no deficits in ability or want to expend effort but inefficiently scale high effort choices with high reward magnitude or probability. Specifically, compared to controls, they have reduced capacity to rely on expected value to optimize effort allocation^45–47^. To investigate this in our model, we ran a series of reward-based operant tasks. Adolescent animals (P35±1 day) first learned to execute a nose poke in response to a light cue to receive a sucrose pellet reward (FR1), then progressed to 2 sessions of executing 5 nose pokes to receive the reward (FR5) (**Figure 3a**). In these low-effort operant sessions, control and DA-*Grin2a* KO animals performed similarly. Cue-action latencies, action-reward latencies, and total completed trials were not significantly different between groups (**Figure 3b-d**), suggesting basic operant reward association learning was conserved in the DA-*Grin2a* KO animals. Animals were then tested on a progressive ratio task for 4 sessions. The task began with an FR1 trial, but in each subsequent trial the response requirement to receive a reward increased exponentially (**Figure 3e**). Progressive ratio allows for measures of motivation through breakpoint (or highest completed ratio) and effort optimization through response rate and post reinforcement pause (PRP) duration^48^. DA-*Grin2a* KO adolescents did not differ from controls in their breakpoint (**Figure 3f**).

**Figure 3:**
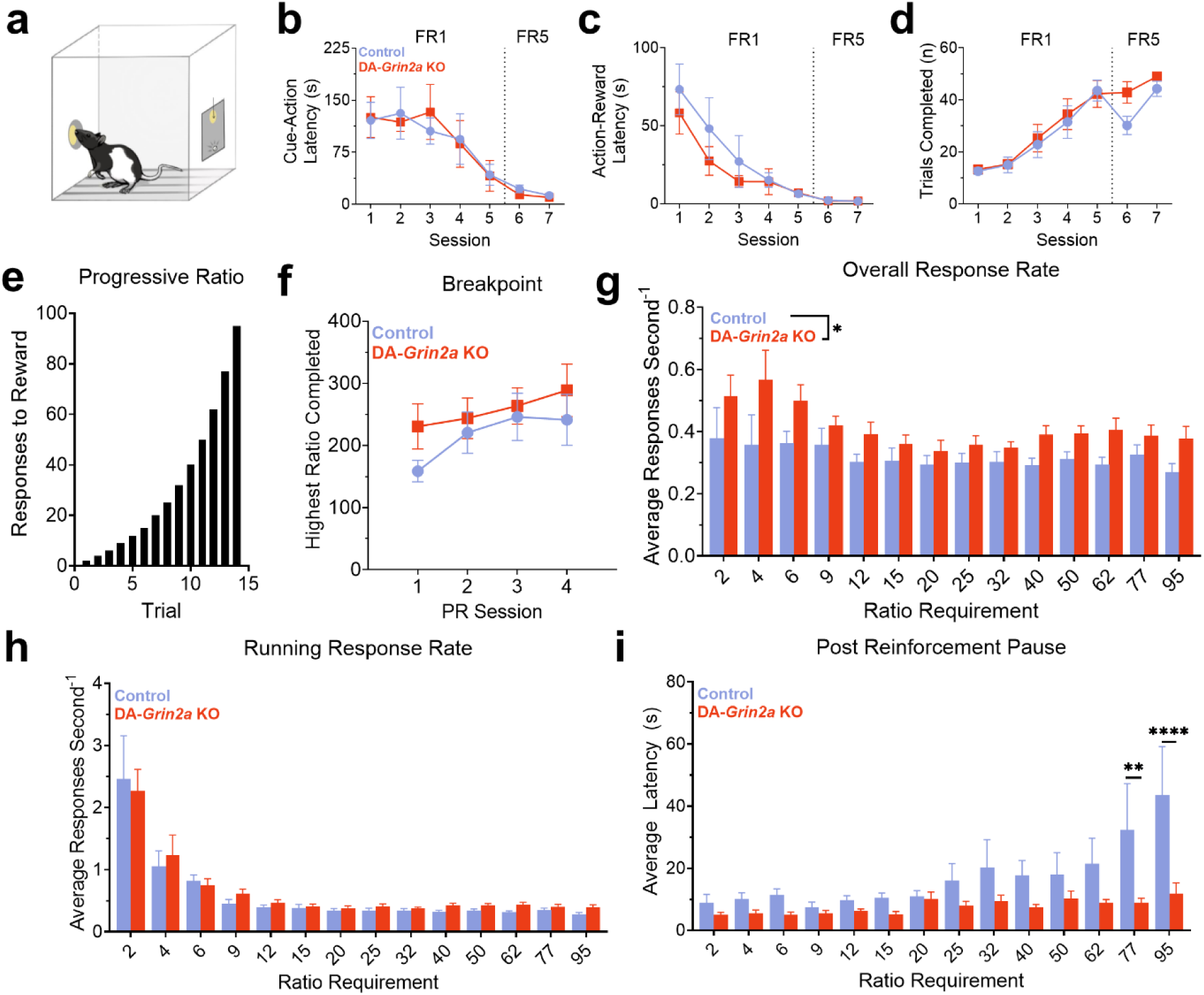
Adolescent DA-*Grin2a* KO results in disrupted effort optimization in progressive ratio. **a**, Representative schematic of rat behaving in operant box during the progressive ratio (PR) task. **b, c, d,** Data collected during FR1 and FR5 training sessions show no group differences in basic operant behavior. Cue-action latency (b), action-reward latency (c), and total trials completed (d) were similar between groups. **e,** Graphic representation of exponential increase in responses required with each trial during the progressive ratio (PR) schedule. **f,** DA-*Grin2*a KO animals showed no differences in breakpoint from controls across the four PR testing days. **g,** DA-*Grin2a* KOs had higher overall response rates compared to control animals, main effect of treatment (*p=0.0335). **h,** No difference between groups in running response rate. **i,** Significant interaction between treatment and ratio requirement (\*\**p*=0.0065) in average post-reinforcement pause (PRP), post hoc analysis revealed significant difference in PRP when ratio requirement was 77 (**p=0.0050) and 95 (****p<0.0001); (control *n*=10, DA-*Grin2a* KO *n*=10 rats). Significance testing: two-way ANOVA: Sidak post hoc. Error bars denote ± SEM.

However, while animals in both treatment groups completed a similar number of trials, their overall response rates significantly differed in these sessions (**Figure 3g**). Overall response rate was calculated as total responses divided by total trial time, so increased overall response rates could reflect either enhanced motor capacity (i.e., faster responding) or increased motivation to engage in effortful behavior (i.e., shorter initiation delays)^48^. To dissociate these contributions, we calculated running response rates (total responses per second following the first active response) and PRPs (latency from cue onset to trial initiation). We observed no difference in running response rates between groups (**Figure 3h**), but a significant difference in PRPs (**Figure 3i**). Typically, PRPs increase as a function of an increasing response requirement^48^ as observed in control animals. The DA-*Grin2a* KO adolescents, however, did not flexibly change this response and maintained stable, and shorter, response latencies despite the increase in action requirement. Thus, while general motivational responding was normal in the DA-*Grin2a* KO animals, their abnormal post reinforcement behavior in high effort trials suggests disrupted effort optimization and impaired ability to use feedback to modify motivated actions.

#### Pavlovian learning and reversal learning

Individuals with SCZ display impairments in reversal learning^49,50^. To test whether this deficit occurs in our adolescent rat model, valence-specific behavioral flexibility was assessed with a flexible contingency learning (FCL) task^51–53^. Adolescent animals (P35±1 day) were exposed to two novel conditioned stimuli (CS), which appeared in random order during the behavior session, each of which predicted the delivery of an unconditioned stimulus (US): either a reward or shock. After 5 sessions, during which associations with each CS were learned, the contingencies were switched so the CS that was initially associated with reward (referred to as CS_A_) now predicted shock and the CS that previously predicted shock (referred to as CS_B_) was now followed by reward delivery (**Figure 4a**). The key behavioral readout for this task was discrimination index (DI), which was the number of reward port entries an animal made during each CS normalized to entries made during the inter-trial-interval (ITI). The reward-predictive CS_A_ and its subsequent reversal to shock-predictive was learned similarly by both DA-*Grin2a* KO and control animals (**Figure 4b**). The shock-predictive CS_B_ was also learned similarly between groups as demonstrated through suppressed DI values. After reversing the CS_B_ association to reward, however, DA-*Grin2a* KO animals developed higher levels of conditioned discrimination compared to controls (**Figure 4c**). Additional analysis of these data showed this difference in DI was not due to faster latencies to respond to the cue and, while not significant, may be more influenced by approach probability (**Figure S2**). To determine if the observed behavioral difference was driven by distinct learning processes, we applied computational models rooted in Pavlovian learning theory to the FCL data. We employed two distinct models: the Rescorla-Wagner model, which infers learning from immediate prediction errors with a fixed learning rate, and the Pearce-Hall model, which predicts learning from stimulus associability based on prior prediction errors and allows for more flexible learning rates. These models both work to explain how associations form between stimuli and outcomes, but the more flexible Pearce-Hall model accounts for the influence of earlier stimuli presentations on information updating and related phenomena such as latent inhibition that are not captured with the Rescorla-Wagner model^54,55^. Models were fit to data collected during the initial 5 reward association learning sessions and fit both control and DA-*Grin2a* KO data with no clear preference for either learning mechanism (**Figure 4d, Figure S3**). Data simulated using individual parameter estimates successfully recapitulated our observed data during acquisition (**Figure 4e**). These similarities were expected because with no prior learning about the reward-predicting stimulus, both models relied on similar principles of associative learning. When the models were applied to data from reward association learning *after* the contingency reversal, we observed that the control group’s behavior was better explained by Pearce-Hall whereas the DA-*Grin2a* KO animals’ behavior was well described by Rescorla-Wagner (**Figure 4f, Figure S3f,h**). Importantly, data simulated with parameter estimates from the preferred model were able to recover the group differences we initially observed (**Figure 4g)**, while the unpreferred models tended to misestimate behavioral trajectories within groups (**Figure S3e,g**). These results indicate that DA-*Grin2a* KO animals expressed a reduced capacity to gate associative processes based on previous learning, and this led to altered behavioral adaptation when outcome contingencies switched.

**Figure 4:**
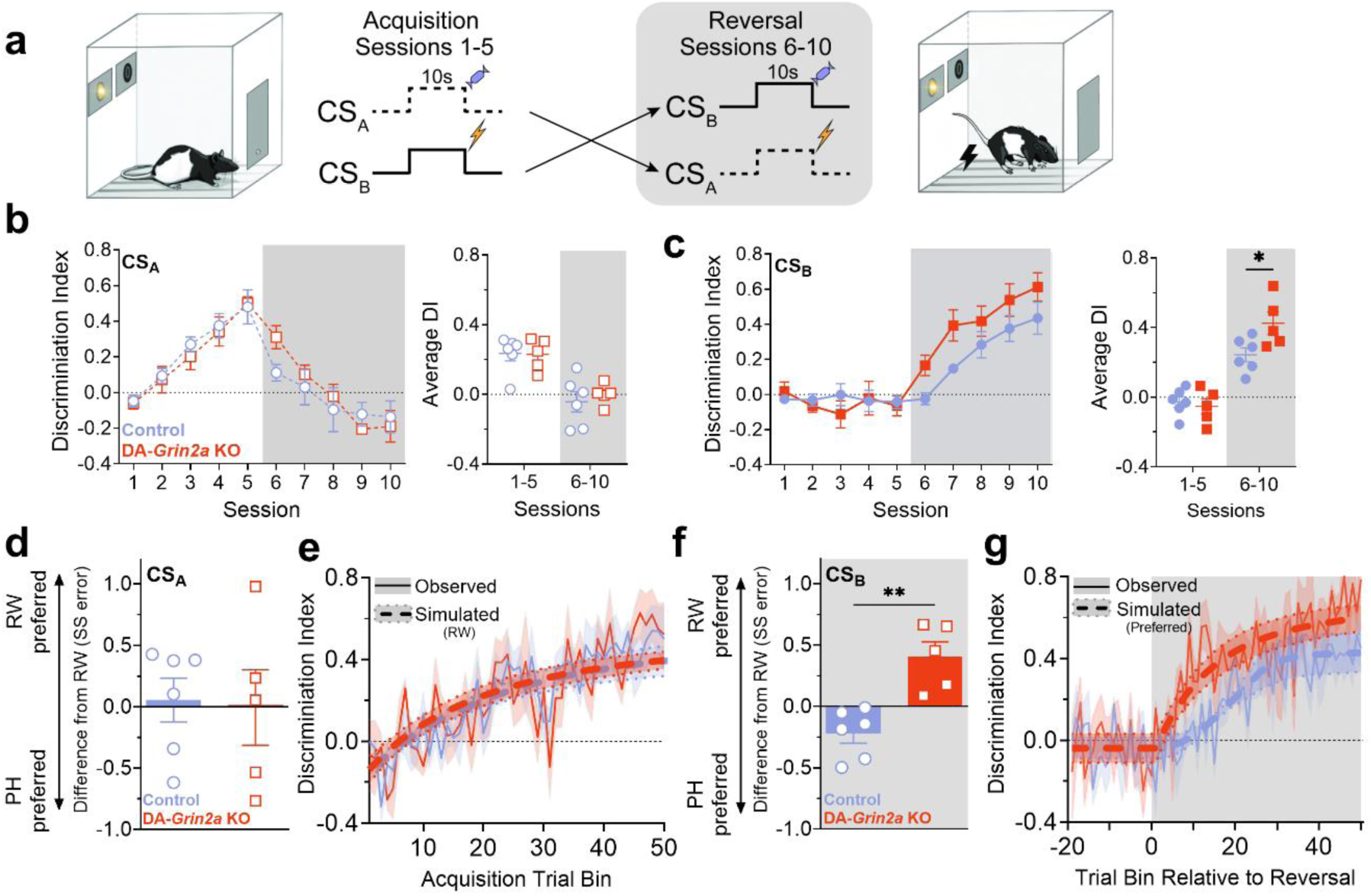
Computational modeling of valence specific behavioral flexibility reveals altered associative learning in DA-*Grin2a* KO adolescents. **a**, Schematic depicting flexible conditioning learning (FCL) task. **b,** Discrimination index (DI) over sessions and averaged across sessions were used as a behavioral readout for the FCL task. Both treatment groups responded to CS_A_ similarly, increasing food port entries relative to baseline during CS_A_ presentation, then decreasing once the CS_A_ contingency was switched on Session 6. **c,** DI over sessions and averaged across sessions in response to CS_B_. There were no significant differences between treatment groups in response to CS_B_ over all sessions, but when DIs were pooled across initial learning (Session 1-5) and reversal (Session 6-10) DA-*Grin2a* KO animals had significantly higher DIs than controls after the contingency switch (\**p*=0.0204). **d,** Models were fit to data from acquisition (Sessions 1-5) of the reward association (CS_A_). Average and individual values (symbols) for difference in model fit between Rescorla Wagner (RW) and Pearce-Hall (PH). Values <0 indicate a better fit from PH mechanisms, >0 indicate a better fit from RW. No model was preferred for the initial learning data. **e,** Observed (solid lines) and simulated (dashed lines) DIs for the simplest (RW) model for each group recovered behavioral trajectories over acquisition across both groups. **f,** Models were fit to data around the contingency reversal (Session 4-10) for CS_B_ (shock to reward association). Average and individual values (symbols) for difference in model fit between RW and PH. The RW model was preferred for all DA-*Grin2a* KO animals, while the PH model was preferred for control animals (\*\**p*=0.0016). **g,** Observed (solid lines) and simulated (dashed lines) DIs for the best fit model for each group recovered differences in behavioral trajectories over reversal between groups; (control *n* = 6, DA-*Grin2a* KO *n* = 5 rats). Significance testing: unpaired t-test. Error bars and shading denote ± SEM.

### DA-Grin2a KO disrupts dopamine release in young adult animals

Disruptions in striatal dopamine neurotransmission are consistently reported in people with SCZ and in individuals at high risk to develop this disorder^24,36,37,56^. Notably, these disruptions are subtle and do not involve overt changes in dopamine related measures in postmortem or imaging studies. Instead, PET and fMRI studies have reported small but consistent disruptions in presynaptic activity^57^ and in prediction error signaling in people with SCZ^58,59^. To determine the long-term consequences of our DA-*Grin2a* KO on prediction error signaling, we measured dopamine in the ventral striatum (VS) of young adult animals (P60+) preforming the FCL task using fiber photometry with a fluorescent dopamine reporter (GRAB_DA_) (**Figure 5a,b**)^60^. This event-rich task allowed us to assess phasic dopamine responses to novel stimuli, their transition to predictive stimuli, as well as to expected and unexpected outcomes. Of note, these measures have two caveats. First, the time constraint of rodent adolescence and other technical considerations did not allow us to run these studies in adolescent animals, when the above behavioral measures were performed. Second, the animals were tethered during the task, potentially influencing their movement in the operant box, which may have contributed to no observed difference in DI in the young adult animals (**Figure S4a**). Given these limitations, we primarily focused on analyses of fiber photometry data that would provide fundamental information about CS processing as opposed to correlation of dopamine signals with behavioral events.

**Figure 5:**
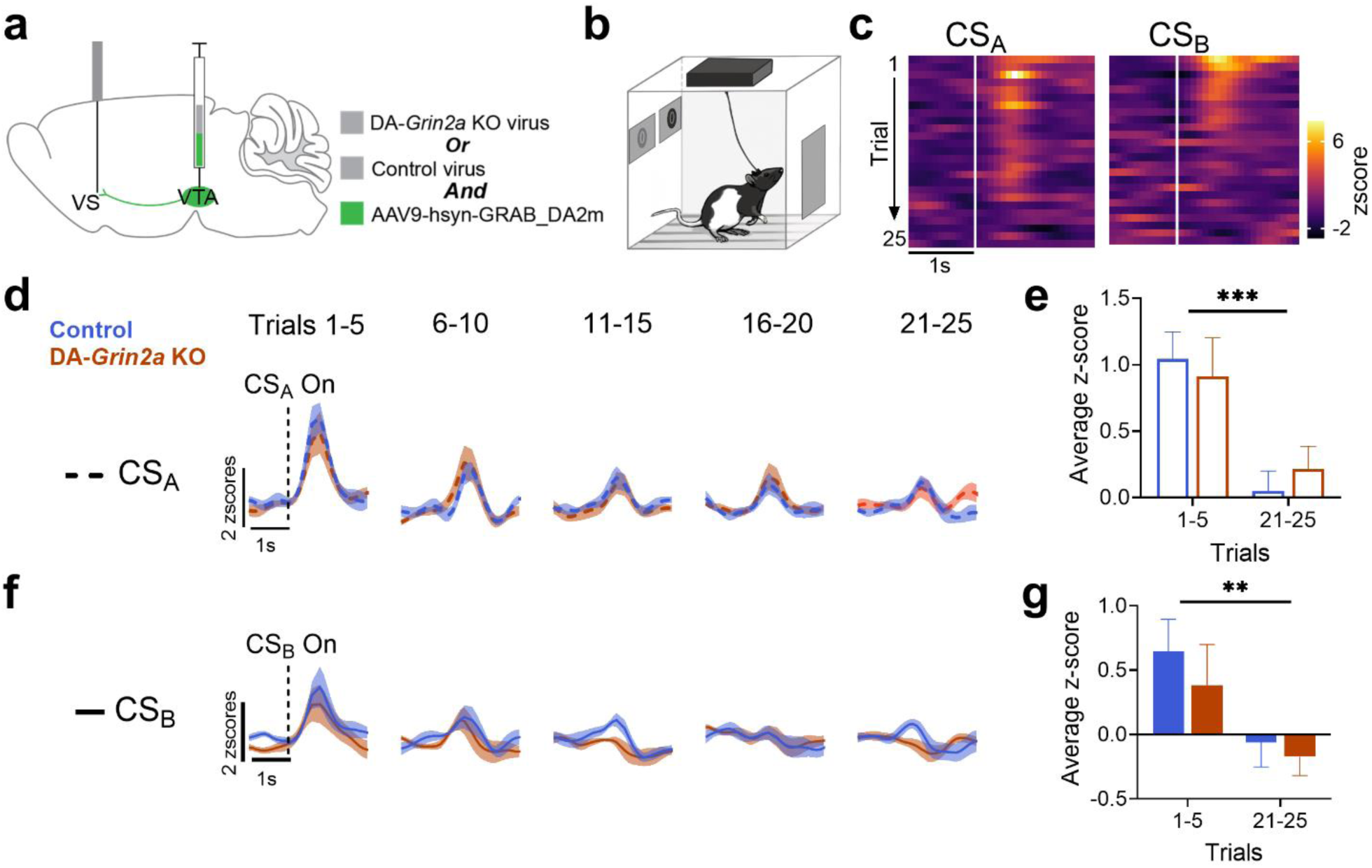
Dopaminergic response to novelty in the ventral striatum is conserved in young adult animals with the DA-*Grin2a* KO virus. **a**, Schematic of virus infusion and fiber implant locations. **b,** Schematic of fiber photometry paradigm in operant box during FCL. **c,** Representative heatmaps from single animal showing first 25 presentations of CS_A_ and CS_B_ during Session 1. **d,** Average fluorescent signal, binned into 5 trial sections for the first 25 presentations of CS_A_ in Session 1. **e,** Average z-score across 2 s following CS_A_ onset showed reduction in response from first 5 presentations to final 5 presentations (\*\*\**p*=0.0003). **f,** Average fluorescent signal for each animal, binned into 5 trial sections for the first 25 presentations of CS_B_ in Session 1. **g,** Average z-score across 2 s following CS_B_ onset showed reduction in response from first 5 presentations to final 5 presentations(\*\**p*=0.0073); (control *n* = 7, DA-*Grin2a* KO *n* = 8 rats). Significance testing: two-way ANOVA. Error bars and shading denote ±SEM.

Dopamine responses to novelty were assessed by quantifying phasic responses to CS_A_ and CS_B_ onset in early trials of Session 1—before any association learning had occurred (**Figure 5c**). To quantify this response, we binned data from the first 25 presentations of CS_A_ into 5 trial bins (**Figure 5d**), then calculated the average z-score across the 2 s following CS_A_ onset. Both groups’ response to CS_A_ decreased over time, and we found no significant differences between control and DA-*Grin2a* KO animals (**Figure 5e**). A similar pattern was observed in response to CS_B_ (**Figure 5f,g**), suggesting novelty signaling was not disrupted by DA-*Grin2a* KO manipulation.

To determine if there were differences in dopaminergic responses once initial associations were learned, we quantified signals during CS onset and US delivery in Session 5 (**Figure 6a, S5b**). Both groups displayed increases in ventral striatal dopamine in response to the reward-predicting cue (CS_A_) and reduced dopamine in response to the punishment-predictive cue (CS_B_) (**Figure 6b**). We observed a similar pattern in response to the two USs, with reward delivery resulting in increased dopamine and a reduced dopamine response following shock delivery (**Figure 6c**). Thus, we observed expected dopamine signaling^61^ in DA-*Grin2a* KOs during initial association learning. We then quantified the dopaminergic response when the cue-outcome contingencies were reversed in Session 6. We split Session 6 into Early trials (first 25 presentations) and Late trials (final 25 presentations) to determine differences as the new associations were formed. During CS_A_ trials where animals were expecting a reward but instead received a shock (**Figure 6d, S5c**), we observed the expected decrease in dopamine signal to CS_A_ onset across Early to Late trials as the cue began to lose its positive association (**Figure 6e**). In response to the unexpected shock associated with CS_A_, control animals showed the expected heightened dopaminergic response in Early trials, which decreased by the Late trials, suggesting this was a response to the mismatch between expected and actual outcome (**Figure 6f**).

**Figure 6:**
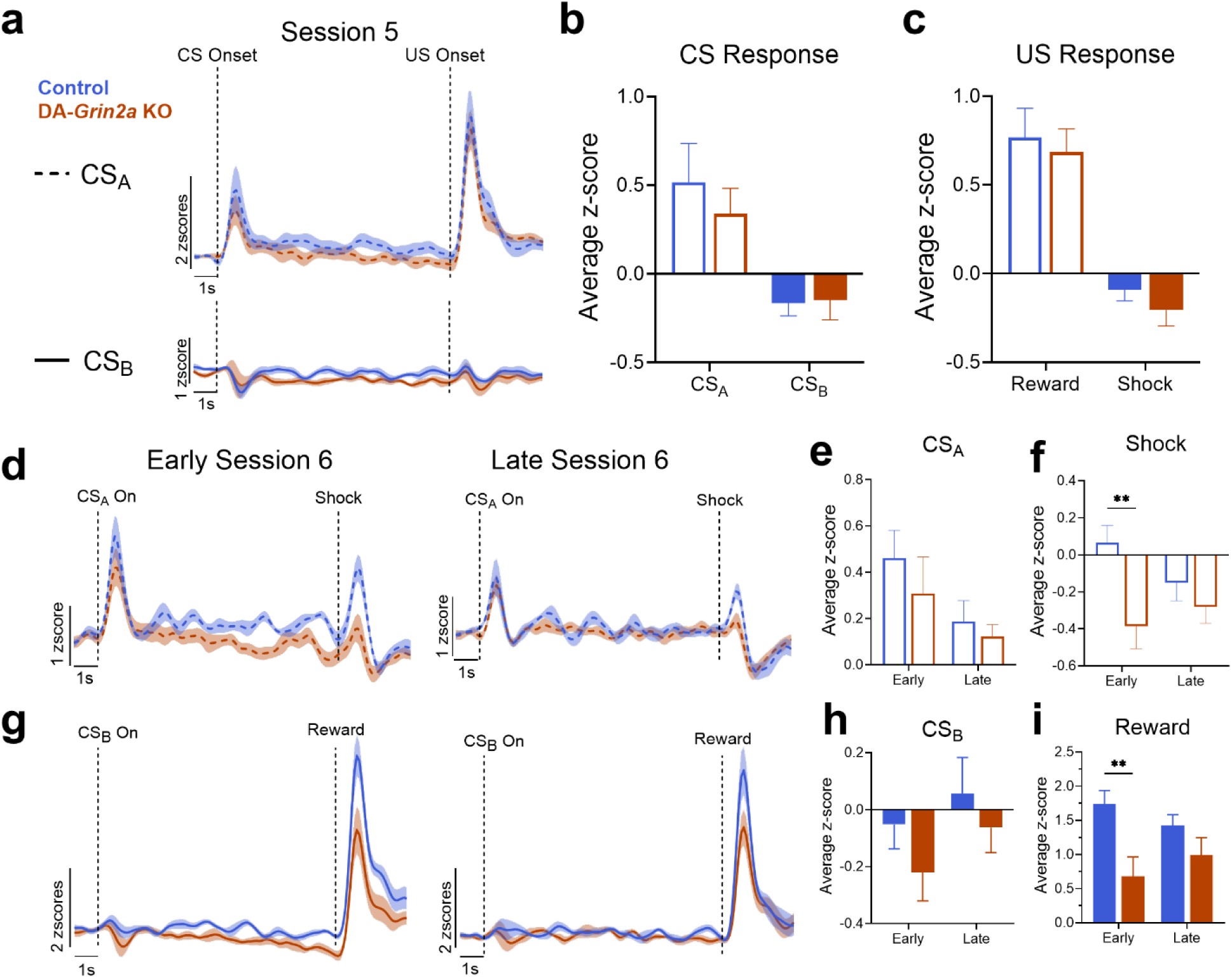
DA-*Grin2a* KO young adults show abnormal dopaminergic response to unexpected outcomes during FCL. **a**, Fluorescent signal of CS_A_ and CS_B_ trials on Session 5 of FCL. **b,** Average z-score across 2 s period at CS delivery showed positive responses to the reward-predicting cue (CS_A_) and negative responses to the shock-predicting cue (CS_B_) in both groups. **c,** Average z-score across 4 s period at US onset showed a similar pattern. **d,** Average fluorescence response during Early and Late CS_A_ trials of Session 6. **e**, Average z-score across 2 s period at CS_A_ onset, no significant group differences. **f,** Average z-score across 4 s period at shock delivery, control animals had significantly higher responses than DA-*Grin2a* KOs in early trials that fell by late trials (\*\**p*=0.0081). **g,** Average fluorescence response during Early and Late CS_B_ trials of Session 6. **h**, Average z-score across 2 s period at CS_B_ onset, no significant group differences. **i,** Average z-score across 4 s period at reward delivery, control animals had significantly higher responses than DA-*Grin2a* KOs in early trials that fell by late trials (\*\**p*=0.0336); (control *n* = 7, DA-*Grin2a* KO *n* = 8 rats). Significance testing: two-way ANOVA: Sidak post hoc. Error bars and shading denote ±SEM.

Interestingly, DA-*Grin2a* KO rats showed a significantly muted response to the unexpected shock during Early Session 6 trials (**Figure 6f**). During CS_B_ trials where the cue that was initially associated with shock switched to reward (**Figure 6g, S5c**), we similarly observed the expected increase in response to CS_B_ onset across Early to Late trials as the cue began to lose its negative association (**Figure 6h**). As with CS_A_ trials, we observed a deficit in dopaminergic response to the unexpected reward in DA-*Grin2a* KOs compared to controls (**Figure 6i**). Together these data suggest that young adult animals with DA-*Grin2a* KO have disrupted dopamine signaling in response to unexpected outcomes, regardless of valence. As previously discussed, the behavioral consequences of this signaling deficit are unclear. Dopamine levels differed between control and DA-*Grin2a* KOs in Early trials of Session 6; however, there were no significant differences between the groups in terms of probability or latency to approach the food port during these trials (**Figure S6**). In later sessions of FCL, control animals showed increased probability to approach during CS_B_ presentation (**Figure S4c**). However, fiber recording data from Session 10 revealed no significant signaling differences between groups at that time (**Figure S7**). These findings suggest that while manipulating *Grin2a* expression led to disruptions in dopaminergic signaling, the effect of these differences on behavior remains unclear under the tested conditions. Future work will explore how this altered dopamine modulation affects behavior in different contexts.

## DISCUSSION

Informed by both GWAS and exome sequencing studies^9–11,62^, we developed a rodent model with construct and face validity that provides insight about how variants of *GRIN2A* can lead to delayed onset of a collection of behavioral and dopamine abnormalities relevant to positive symptoms of SCZ. Our approach was guided by the well-established clinical literature showing a relationship between dopamine neurotransmission and psychosis^24,25,36–38,56,63–65^, which often emerges later in development than cognitive and affective symptoms. *GRIN2A* encodes the GluN2A subunit of the NMDA receptor and has a cortical developmental expression profile that corresponds with the emergence of cognitive symptoms in childhood and early adolescence. We found that—unlike cortical regions where GluN2A levels remain stable throughout adolescence—there was significant loss of GluN2A in the dopamine neuron-containing regions of the midbrain during this developmental period. This led us to reason that the *GRIN2A* variants observed in the clinical population may exacerbate this developmental decline and contribute to the emergence of psychosis. We find that a genetically driven deficit of *Grin2a* in midbrain dopamine neurons of adolescent rats is sufficient to recapitulate some behaviors that are consistent with psychosis. These findings provide an etiologically relevant model for latent expression of psychosis which may aid with the development of novel course alternating treatments for SCZ.

### *GRIN2A* and schizophrenia

Genetic variants in *GRIN2A* have been associated with SCZ diagnosis^9–11,62^, providing further credence to the long-standing theory that abnormalities in NMDA receptor function are a core component of SCZ pathophysiology^1,66^. Global heterozygous loss of *Grin2a* in mice produces sustained phenotypes such as motor abnormalities, disrupted cortical activity, and changes in expression of multiple genes relevant to SCZ^15^. These global KO studies prompted our investigation into the consequences of an age- and cell type-specific *Grin2a* manipulation. We were particularly interested in how a *GRIN2A* loss of function may lead to latent expression of psychosis and dopamine processing deficits^24,25,35–38^. This led us to ask if selective loss of function of *Grin2a* in the dopamine neurons of adolescents is sufficient to produce phenotypes that are relevant to psychosis. We used a Cre-driven CRISPR/Cas9 virus in adolescent Th::Cre male and female rats to induce a deficit in *Grin2a* in midbrain dopamine neurons. This approach was successfully implemented despite the short duration of rodent adolescence. Moreover, the animals lacked any gross motor or learning deficits as measured by operant, Pavlovian, and maze tasks. The behavioral and dopamine release changes we observed were highly selective to constructs relevant to psychosis including hypersensitivity to psychostimulants, inefficient effort optimization, and reduced capacity to utilize feedback to guide behavior.

### DA-*Grin2a* KO results in adolescent phenotype relevant to positive SCZ symptoms

Clinically, SCZ is defined as a heterogeneous psychiatric disorder characterized by the presence of positive, negative, and cognitive symptoms. To determine the extent of the effect of our manipulation, we included a variety of SCZ-related behavioral tasks in our battery. We found a specific behavioral phenotype in which adolescent animals with the DA-*Grin2a* KO showed abnormalities in line with early emerging positive symptoms.

Consistent with clinical studies showing the exacerbation of positive symptoms in people with SCZ by psychostimulants, we determined DA-*Grin2a* KO adolescents had increased sensitivity to the locomotor effects of amphetamine. When effort-based reward learning was assessed with an exponential progressive ratio, we observed control rats suppressed their stimulus-driven responding in high effort trials, while DA-*Grin2a* KO animals did not. In the classical conditioning task, behavioral differences between control and DA-*Grin2a* KO animals were observed selectively during contingency reversal sessions. These behavioral data, reinforced with modeling, revealed DA-*Grin2a* KO animals showed reduced capacity to use feedback to guide behavior by incorporating previous learning with new experiences. Together these data reflect clinically observed phenomena relevant to positive SCZ symptoms.

### DA-*Grin2a* KO results in disrupted dopamine signaling

Abnormalities in dopamine neurotransmission in people with SCZ are consistently reported. Evidence for faulty prediction error signaling in schizophrenia comes from fMRI studies showing abnormalities in regional activity during reinforcement learning.

Specifically, people with SCZ have reduced activity in the midbrain and ventral striatum when presented with an unexpected outcome, while maintaining a typical level of activity in response to expected outcomes^67,68^.

To drive learning, dopamine neurons increase activity when an unexpected event occurs^69,70^. After the contingency reversal in the FCL, the robust increases in synaptic dopamine in Early trials observed in control animals was attenuated in the DA-*Grin2a* KOs. This suggests that the DA-*Grin2a* KO selectively impairs the ability of dopamine neurons to increase signaling when there is a difference between expected vs actual outcomes. These data demonstrate a specific role for GluN2A-containing NMDA receptors on midbrain dopamine neurons in prediction error signaling in response to unexpected stimuli. This is consistent with clinical data^67,68^, providing a potential mechanism through which a genetic variant of *GRIN2A* could contribute to the deficits in prediction error encoding observed in people with SCZ.

### Conclusion and Limitations

Our findings provide mechanistic insight into how variants of *GRIN2A* may lead to the latent presentation of abnormalities in dopamine dynamics in SCZ. Our aim was not to produce a full model of *GRIN2A* loss of function, but to tease apart how specific cell-types and neural systems may be affected by this mutation. While the mutation is not confined to a single region in humans, our approach is mechanistically informative and provides a novel potential model for designing and testing course-altering treatments for transition to SCZ in vulnerable individuals. Decades of translational studies focused on the NMDA receptor have led to the discovery of potential treatments for SCZ^1,71,72^, some of which showed clinical efficacy for symptom amelioration only in younger individuals^73^. Clinical trials with these drugs may have failed in the past because they were tested in chronically ill patients with decades of antipsychotic drug treatment, which may have remodeled and changed the nature of dopamine abnormalities. In younger people with—or at high risk to develop— SCZ, these targets or newer GluN2A-selective positive allosteric modulators may provide a precise approach for preventing or altering the course of the illness.

## METHODS

### Subjects

All animal procedures were approved by the Institutional Animal Care and Use Committee at Oregon Health & Science University. Long Evan’s male and female rats were bred in house and maintained on a reverse 12hr light/dark cycle with ad libitum access to food and water unless otherwise stated.

### Tissue collection and protein extraction

Brain levels of GluN2A were examined in animals aged P21, P35, P45, and P80. Tissue collection methods were adapted from Wager-Miller^74^. Briefly, at each age of interest, brains were harvested, and flash frozen in isopentane that was pre-chilled with dry ice. Whole brains were stored in conical tubes at -80°C until dissection.

While maintaining a frozen condition, brains were sliced into 1mm sections using a pre-chilled brain matrix and razor blades. Bilateral micropunches of tissue were collected from regions of interest using a rat brain atlas as a reference^75^. To extract protein, tissue samples were homogenized in a lysis buffer containing protease and phosphatase inhibitors. Protein concentration was quantified with a Pierce BCA Protein Assay Kit (ThermoFisher). Homogenates were aliquoted and stored at -80°C until use.

### Protein quantification

Automated immuno-quantification was run on a Wes instrument (Bio-Techne) according to the manufacturer’s protocol^76^. Briefly, each tissue sample was diluted to a concentration of 0.08 μg/μL protein and mixed with dithiothreitol and a fluorescent molecular weight marker at a ratio of 5:1. These samples were heated to 95°C for 5 min for protein denaturation and then loaded into the allocated wells of the Wes plate. Other wells were loaded with blocking reagent, primary antibody (Rabbit-anti-NMDAR2A (Sigma Aldrich, #M264)), horseradish peroxidase (HRP)-conjugated secondary antibody, and a mix of Luminol-S and Peroxide (Bio-Techne Anti-Rabbit Detection Module, #DM-001). Size-based separation electrophoresis, immobilization, and immunodetection were automatically run using the capillary array system of the Wes (**Figure S1**). All samples were normalized to total protein as a loading control (Bio-Techne Total Protein Detection Module for Chemiluminescence, #DM-TP01). Densitometric analysis was performed using the Compass software (v5.0.1) from Bio-Techne.

Tissue dissection, micropunching, protein extraction, and Western blots were run as described above on punches collected from the VTA of virus infused animals.

### Viral production

The Cre-inducible recombinant adeno-associated virus (AAV) vector constructs utilized in this study were produced and gifted by Larry Zweifel^77^. Viruses in these experiments were: AAV1-FLEX-Cas9-U6-sgGrin2a for dopamine neuron-specific *Grin2a* knockout (DA-*Grin2a* KO) (Addgene, #124850) which targets the first conserved exon (exon 6) across splice variants in both rat and mouse (sgRNA: 5’ GAGCAGGCAACCGGCTTGCCC 3’). AAV1-FLEX-Cas9-U6-sgRosa26 was used as a control (Addgene, #159914) (sgRNA 5’ CTCGATGGAAAATACTCCGAG 3’). These viruses were co-infused with AAV1-hSyn-DIO-EGFP (Addgene, #50457) at a 3:1 ratio to aid with visualization in subsequent experiments.

### Stereotaxic surgeries

Juvenile rats (P21±1), were anesthetized with isoflurane and placed in a stereotaxic apparatus and body temperature was maintained using a water heating circulation pump (E-Z Systems). The scalp was incised and two bilateral craniotomies over the VTA were performed. Four 250 nL infusions at the bilateral VTA sites (AP -4.9 mm, ML ±0.7 mm, DV - 6.5 and -6.0 mm from dura) were delivered using a syringe pump (World Instruments) and micro infusion syringe (Hamilton) at a rate of 100 nL/min. To prevent backflow, the syringe was not removed until 5 min after infusions. The incision was then closed with surgical sutures, treated with triple antibiotic, and animals were administered 5 mg/kg carprofen subcutaneously to relieve pain and inflammation. Animals remained on heat and were administered oxygen until consciousness was regained, then returned to their home cage. There was a minimum of a two-week incubation period before experiments were run.

For fiber photometry experiments, juvenile (P21±1) animals were infused with either the control or DA-*Grin2a* KO virus mixed with AAV9-hsyn-GRAB_DA2m (GRAB_DA_) (addgene, #140553) at a 1:1 ratio. These were infused bilaterally into the VTA as described above.

Once animals reached adulthood (P60-70), they were implanted with a 400 µm optical fiber (Doric Lenses Inc) into the ventral striatum (AP +1.6 mm, ML 1.5 mm, DV -7.0 mm). Animals were given at least one week to recover from the implant surgery before behavioral training.

### Immunohistochemistry

Animals were euthanized with an intraperitoneal (IP) injection of 400 mg/kg chloral hydrate and perfused via the vascular system with 0.1 M phosphate buffer solution (PBS) followed by 4% paraformaldehyde (PFA). Brains were removed and stored in 4% PFA at 4°C overnight, then transferred to a 30% sucrose solution and stored at 4°C until sectioning.

Coronal sections (35 μm) of each brain were collected using a cryostat (Lecia) and stored in 0.1 M PBS with 0.05% sodium azide. Free-floating sections were blocked and permeabilized at room temperature for 2 hrs in 0.1 M PBS, 0.25% Triton X, 10% normal donkey serum. Sections were then incubated overnight at 4°C with the following primary antibodies: chicken anti-TH (1:500, #76442, abcam), rabbit anti-GFP (1: 500, #290, abcam), or mouse anti-HA (1:500, MMS-101P, Biolegend). Sections were washed, then incubated in secondary antibodies (AlexaFluor-594; 488, 1:1000) for 2 hrs at room temperature. Sections were washed again and mounted onto Super Frost+ slides.

Coverslips were applied using Vectashield HardSet antifade mounting media. Fluorescence was imaged using a Ziess Axiovert 200 microscope and cell counting was done in Zen 2 software (Zeiss).

For fiber photometry experiments, sections containing the fiber tract were mounted onto Super Frost+ slides. Coverslips were applied using Vectashield HardSet antifade mounting media containing DAPI and imaged using a Ziess Axiovert 200 microscope and cell counting was done in Zen 2 software (Zeiss). To confirm fiber placement in the ventral striatum, images were superimposed with a brain atlas^75^ (See hit map, **Figure S4a**).

### Slice Electrophysiology

Two to three weeks following virus infusion (P35-42), rats were anesthetized with isoflurane and decapitated. The brain was collected and deposited in artificial cerebrospinal fluid (ACSF) at 32–35°C containing (in mM): 126 NaCl, 2.5 KCl, 1.2 MgCl_2_, 2.4 CaCl_2_, 1.4 NaH_2_PO_4_, 25 NaHCO_3_, and 11 Dextrose. The ACSF used for brain extraction, slicing and recovery also contained 2 mM kynurenic acid (Sigma) to prevent NMDA-mediated excitotoxic damage. Horizontal slices (220 μm) containing the midbrain were cut using a vibratome (Leica) in warm ACSF bubbled with 95% O_2_/5% CO_2_. Slices were allowed to recover in the same buffer at 30°C for at least 30 min prior to recording. Hemisected slices were transferred to the recording chamber, which was continuously perfused at 2 to 3 mL/min with bubbled ACSF at 34°C. In most experiments the recording buffer was identical to the collection buffer except it did not contain any MgCl_2_, to allow for NMDA receptor activation by glutamate iontophoresis. In a subset of recordings, experiments were started in Mg-containing ACSF and Mg-free ACSF was perfused later. All experiments were completed within 7 h of slicing.

VTA dopamine neurons were identified by their morphology and the presence of GFP. Recordings were made using glass pipettes (1.0 – 2.5 MΩ resistance) filled with an internal solution containing (in mM): 100 K-gluconate, 20 NaCl, 1.5 MgCl_2_, 10 HEPES (K), 2 ATP, 0.2 GTP, 10 phosphocreatine, and 10 BAPTA (4K). Recordings were made in whole-cell configuration. After break-in, cells were voltage-clamped at −55 mV and resistance and capacitance were monitored; cells with a series resistance ≥12 MΩ were discarded. The dopaminergic identity of the patched cells was further confirmed by the presence of a hyperpolarization-induced depolarizing current (I_h_). Cells with an I_h_ < 200 pA were discarded. Another glass pipette was filled with a 1 M solution of monosodium glutamate (Sigma Aldrich). This pipette was lowered into the slice and placed in proximity to the patched cell, while a continuous backing current of +1.0 nA was applied. Glutamate was iontophoretically delivered by the application of 20 ms negative current pulses (-40.0 to - 80.0 nA). Location and current intensity were adjusted to produce a stable depolarizing current in the patched cell.

Data were acquired with AxoGraph software (Berkeley, CA) and recordings were monitored with LabChart (AD instruments, Colorado Springs, CO). Current responses in Mg-free ACSF were recorded until at least 10 stable responses were obtained. For experiments where MG-free ACSF was perfused after patching this took up to 25 min. The slice was then perfused with Mg-free ACSF containing the GluN2A-specific antagonist TCN201 (30 μM).

Responses to glutamate puffs were then recorded until a stable state was obtained or for a maximum of 20 minutes. The slice was then perfused with Mg-free ACSF containing 10 μM ifenprodil, a GluN2B-specific antagonist. Responses were measured until a stable state was reached, or a maximum of 20 min. Lastly, we bath applied 50 μM AP5 (Sigma Aldrich), a non-selective NMDA blocker, to confirm the identity of the measured currents. Because both TCN201 and ifenprodil do not readily wash out, each slice was only used for one attempted recording. Recordings were analyzed post-hoc using AxoGraph by averaging the last 5 stable pulses for each condition and measuring the peak amplitude of the resulting trace. Data are expressed as a fraction of the Mg-free baseline.

### Spontaneous behaviors

All behavioral testing was administered at P45-P55. Animals were habituated to handling and the testing environment for two days preceding behavior sessions. All behaviors were run in a dim, white light. On the day of testing, animals were habituated to the room for 2 hr before trials were run. The assays were run as follows:

### Elevated plus maze (EPM)

The EPM consisted of two opposing open arms and two opposing arms bordered with 48 cm walls. Animals were placed in the center of the maze and allowed to explore for 10 min and their movement tracked with PanLab SMART v3.0 software (Harvard Apparatus). Percent of time spent in open and closed arms was evaluated.

### Spontaneous alternation test

A four-arm, plus maze was used to assess spontaneous alternation as a measure of spatial working memory. In this task, distinct visual cues were positioned on all four walls of the behavioral testing room. Animals were placed in the center of the maze and allowed to explore for 10 min. The order of arm entries was recorded manually during the task. Percent alternation and total number of arm entries were evaluated. An alteration consisted of four distinct arm choices out of five consecutive arm entries. A 4/5 alternation score was determined by dividing the total number of alternations into overlapping quintuplets by the number of possible alternations and multiplying by 100.

### Open field (OF) with Pharmacological Challenge

Animals were placed in an opaque square arena and allowed to explore for 20 min. Movement was tracked with PanLab SMART v3.0 software. During this time, baseline locomotion and percent time spent in center (defined as a 5 cm distance from the walls of the arena) was calculated.

After OF habituation, animals received IP injections of 1.0 mg/kg amphetamine (AMPH) and placed back into the arena for 1 hr while video recordings were collected. Horizontal locomotion in cm was tracked with PanLab SMART v3.0 software as animals explored the maze.

### Operant and Pavlovian conditioning

Adolescent behavioral training experiments were performed on animals aged P35-P45. Rats were mildly food restricted and habituated to operant boxes (Coulbourn Instruments) two days prior to experimentation. All behavioral tasks were run in red light and behavior recorded with Graphic State software (Coulbourn Instruments). Each animal received one of the following tasks:

### Progressive ratio of reinforcement

Operant chambers contained one wall with a nose poke hole that could be illuminated and an opposing wall with a food port where sucrose pellets were dispensed. Animals were initially trained on a fixed ratio one (FR1) schedule where one nose poke in response to a light cue resulted in the delivery of one sucrose pellet. After five days of training, animals advanced to an FR5 schedule for two days. For the final four days, animals were tested with a progressive ratio schedule, where the response ratio increased according to the formula 5e(0.2n)^-5^, where n = trial number^78^. This resulted in an exponential increase in response requirement (1, 2, 4, 6, 9, 12 etc.) when rounded to the nearest whole number.

Sessions were terminated if there was a 5 min cessation in responding, if a trial took longer than 45 min to complete, or after 3 hr total. Latencies to respond to the cue, to retrieve a reward, and total rewards received were calculated during FR1 and FR5 sessions. We determined the breakpoint for each animal across the four progressive ratio sessions. For each trial, the overall response rate was calculated as total responses divided by *total* trial time, while running response rate was calculated as total responses per second following trial initiation. Post-reinforcement pause (PRP) duration was defined as the interval from cue onset to the initiation of the next trial after the preceding trial’s completion. Overall response rate, running response rate, PRP were calculated as an average of the testing days. For trial-by-trial analyses in progressive ratio, only trials where 80% of animals completed a given ratio were analyzed.

### Flexible contingency learning (FCL)

Operant chambers contained one wall with a light cue and speaker and an opposing wall with a food port where sucrose pellets were dispensed. We utilized a modified version of a previously published task^51–53^ abbreviated to better fit within the adolescent time window. Briefly, animals were presented with two conditioned stimuli (CS): a 10 sec light or tone. In a counterbalanced fashion, animals were conditioned to pair one CS with the delivery of a sucrose pellet and the other with a mild foot shock (180 ms, 0.2 mA). These unconditioned stimuli (US) immediately followed the termination of either CS. After five days of conditioning, the initial associations were reversed for Sessions 6-10. In each session, animals underwent 100 stimulus pairings (50 trials of each CS) delivered randomly with a 20 sec inter-trial interval (ITI). CS_A_ represents the cue that was initially appetitive and then became aversive, CS_B_ represents the cue that was initially aversive and then switched to appetitive.

Behavior was analyzed by normalizing the amount of nose pokes into the food port during CS to baseline food port entries during the ITI. The following equation was used to calculate this discrimination index (DI):

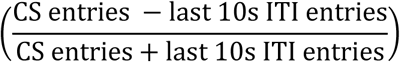

With this formula, a positive DI suggests conditioned approach, a null DI suggests no conditioning has occurred, and a negative DI suggests conditioned suppression. Average DIs across Sessions 1-5 were considered initial learning, while average DIs across Sessions 6-10 were considered contingency reversal learning.

Additional analyses included probability of approaching the food port during the 10 s CS presentation and 10 s following US delivery as well as average latency to approach the food port following CS or US onset.

### Computational modeling

We utilized computational models of learning to further dissect the pattern of responding in individual subjects in each group as measured by the DI during the FCL task (see above).

### Models

We compared two models of associative learning that explain learning as a function of immediate prediction error (Rescorla-Wagner, RW)^79^ or stimulus associability (Pearce-Hall, PH)^80^. In each model, the conditioned strength (V) of a stimulus evolves trial-by-trial according to experience; we predicted the DI as a proxy for the conditioned strength (V) of the stimulus presented on each trial.

RW – Learning in the RW model depends on prediction error (δ) and a fixed learning rate. V develops for each trial according to the difference between actual (λ_n_) and expected outcome (V_n_) for a stimulus on a given trial (n). This prediction error is multiplied by a fixed learning rate (α, free parameter constrained between 0-1) which controls the rate of learning. This can be formulated as:

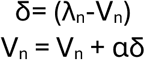

PH - The PH model accounts for learning as a function of the associability (α) of a stimulus. We used an extended formulation of the PH model with gradual learning^81^ where α is updated on each trial from its previous level by a prediction error (δ) that tracks the absolute value of the difference between actual (λ_n-1_) and expected outcomes (V_n-1_) for that stimulus on the prior trial (n-1). This update to the associability of the stimulus is controlled by a free weighting parameter (γ, free parameter constrained between 0-1). The update to V is formulated as:

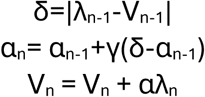

### Model fitting

To address the high trial-by-trial response variability, we smoothed responses using a 5-trial moving average and fit each of the RW and PH models to the DI values for the acquisition (Sessions 1-5) and reversal (Sessions 6-10) epochs for each subject. Initial conditioning strength (V_0_) and value of an outcome (λ) was determined for each subject to account for subject differences in initial bias and outcome sensitivity. Thus, the initial conditioning strength was determined by taking the average DI for the first half of Session 1 (for acquisition) or from pre-reversal (i.e. Sessions 4-5). Value of an outcome was determined by the mean DI observed in the last session of acquisition or reversal. This effectively normalized the initial and maximum ‘strength’ of the cue, leaving the learning trajectory to be fit. Best-fitting model parameters (α and γ for RW and PH, respectively) were estimated through grid search across a range of 1,000 equally spaced values and selected via a least-squares approach.

### Model comparison

Models were assessed by the total sum of squared residual error (SS) and models with the lowest total SS were selected for each group. We also assessed our model selection with a paired permutation approach. Briefly a null distribution was generated by shuffling condition labels within subject 10,000 times (i.e. the within group paired nature of the data was kept intact) and calculating the mean difference between PH and RW residual error.

We then assessed the proportion of instances where a value in the null distribution was equal to or greater than the observed PH/RW difference from the unshuffled data to obtain a p-value (one-tailed). To compare between groups, we quantified the difference in residual sum of squares between RW and PH fits for each subject and compared groups using a two-tailed t-test.

### Data simulations

We validated parameter estimates for α or γ by simulating learning trajectories for each subject from their best fit parameters. We then plotted the group-averaged response of the preferred model to validate the model(s) could recover key trends in the data.

### Statistical analysis

Statistical analyses were performed in GraphPad Prism (Version 10.0.1). Unpaired t-tests were run in instances where two groups were compared. Otherwise, the data were analyzed with either a one- or two-way ANOVA with Tukey or Sidak post hoc corrections, respectively.

### Fiber photometry recording

Two days preceding fiber photometry experiments, implanted animals were mildly food restricted and habituated to handling, being attached to the recording patch cord, and to the testing environment. Animals were then run on the FCL task as described above. Fiber photometry experiments were executed using an RZ10X fiber photometry system (Tucker-Davis Technologies). The Quick-Release Interconnect (Thorlabs) was used to mate the implanted fiber optic cannula and the patch cord. Excitation light was passed through a 400 µm patch cord from 465 and 405 nm LEDs (Doric Lenses Inc), sinusoidally modulated at 220 and 310-Hz, respectively. Signals were demodulated at 1kHz with a 6Hz low pass filter in real-time using Synapse software (Tucker-Davis Technologies). Time-locked behavioral events from the operant box were recorded through the RZ10X digital input ports. Recording was performed on each day of the behavioral task.

### Data analysis

Fiber photometry data were analyzed using Python 3 and the TDT Python package. Raw fluorescence signals from 465 nm (GRAB_DA_) and 405 nm (isosbestic) channels were segmented into trials, each 30 s (10 s before CS onset to 10 s after US onset). For each trial, the GRAB_DA_ signal was fit to the isosbestic signal using a least-squares linear fit (numpy polyfit) to correct for motion artifacts and photobleaching. The fitted signal was used to compute the change in fluorescence (ΔF/F) for each trial.

To assess event-related changes in striatal dopamine, peri-event z-scores were calculated by normalizing the ΔF/F signal to the 10 s baseline period prior to CS onset (within the ITI). Average responses were then computed across trials for each animal. Dopamine responses to specific events were calculated as follows:

Novelty: First 25 CS_A_ presentations and first 25 CS_B_ presentations were grouped into 5 bins of 5 trials each.

Initial association: Responses were averaged across all 50 CS_A_ trials and 50 CS_B_ trials in Session 5.

Association reversal: following reversal in Session 6, responses to unexpected outcomes were assessed by averaging the first 25 trials (Early) and the last 25 trials (Late) of each CS- US pairing.

To quantify differences in synaptic dopamine between the treatment groups, we determined the average z-score across two periods. The CS time bin consisted of the first 2 s following cue onset and US time bin consisted of the first 4 s following the onset of the reward/shock.

Statistical analyses were performed in GraphPad Prism (Version 10.0.1). Average z-scores were analyzed with a two-way ANOVA (with factors Session and Treatment) with Sidak post hoc corrections if there was a significant interaction (p<0.05).

## REFERENCES

1. Moghaddam, B., and Javitt, D. (2012). From revolution to evolution: the glutamate hypothesis of schizophrenia and its implication for treatment. Neuropsychopharmacology 37, 4–15. 10.1038/npp.2011.181.

2. Coyle, J.T., Ruzicka, W.B., and Balu, D.T. (2020). Fifty Years of Research on Schizophrenia: The Ascendance of the Glutamatergic Synapse. Am J Psychiatry 177, 1119–1128. 10.1176/appi.ajp.2020.20101481.

3. Law, A.J., and Deakin, J.F. (2001). Asymmetrical reductions of hippocampal NMDAR1 glutamate receptor mRNA in the psychoses. Neuroreport 12, 2971–2974. 10.1097/00001756-200109170-00043.

4. Weickert, C.S., Fung, S.J., Catts, V.S., Schofield, P.R., Allen, K.M., Moore, L.T., Newell, K.A., Pellen, D., Huang, X.F., Catts, S.V., and Weickert, T.W. (2013). Molecular evidence of N-methyl-D-aspartate receptor hypofunction in schizophrenia. Mol Psychiatry 18, 1185–1192. 10.1038/mp.2012.137.

5. Meador-Woodruff, J.H., Clinton, S.M., Beneyto, M., and McCullumsmith, R.E. (2003). Molecular abnormalities of the glutamate synapse in the thalamus in schizophrenia. Ann N Y Acad Sci 1003, 75–93. 10.1196/annals.1300.005.

6. Javitt, D.C., and Zukin, S.R. (1991). Recent advances in the phencyclidine model of schizophrenia. Am J Psychiatry 148, 1301–1308. 10.1176/ajp.148.10.1301.

7. Moghaddam, B., and Krystal, J.H. (2012). Capturing the angel in "angel dust": twenty years of translational neuroscience studies of NMDA receptor antagonists in animals and humans. Schizophr Bull 38, 942–949. 10.1093/schbul/sbs075.

8. Singh, T., Poterba, T., Curtis, D., Akil, H., Al Eissa, M., Barchas, J.D., Bass, N., Bigdeli, T.B., Breen, G., Bromet, E.J., et al. (2022). Rare coding variants in ten genes confer substantial risk for schizophrenia. Nature 604, 509–516. 10.1038/s41586-022-04556-w.

9. Schizophrenia Working Group of the Psychiatric Genomics, C. (2014). Biological insights from 108 schizophrenia-associated genetic loci. Nature 511, 421–427. 10.1038/nature13595.

10. Ripke, S., O’Dushlaine, C., Chambert, K., Moran, J.L., Kahler, A.K., Akterin, S., Bergen, S.E., Collins, A.L., Crowley, J.J., Fromer, M., et al. (2013). Genome-wide association analysis identifies 13 new risk loci for schizophrenia. Nat Genet 45, 1150–1159. 10.1038/ng.2742.

11. Stefansson, H., Ophoff, R.A., Steinberg, S., Andreassen, O.A., Cichon, S., Rujescu, D., Werge, T., Pietilainen, O.P., Mors, O., Mortensen, P.B., et al. (2009). Common variants conferring risk of schizophrenia. Nature 460, 744–747. 10.1038/nature08186.

12. Trubetskoy, V., Pardinas, A.F., Qi, T., Panagiotaropoulou, G., Awasthi, S., Bigdeli, T.B., Bryois, J., Chen, C.Y., Dennison, C.A., Hall, L.S., et al. (2022). Mapping genomic loci implicates genes and synaptic biology in schizophrenia. Nature 604, 502–508. 10.1038/s41586-022-04434-5.

13. Hall, J., and Bray, N.J. (2022). Schizophrenia Genomics: Convergence on Synaptic Development, Adult Synaptic Plasticity, or Both? Biol Psychiatry 91, 709–717. 10.1016/j.biopsych.2021.10.018.

14. Farsi, Z., Nicolella, A., Simmons, S.K., Aryal, S., Shepard, N., Brenner, K., Lin, S., Herzog, L., Moran, S.P., Stalnaker, K.J., et al. (2023). Brain-region-specific changes in neurons and glia and dysregulation of dopamine signaling in Grin2a mutant mice. Neuron 111, 3378–3396 e3379. 10.1016/j.neuron.2023.08.004.

15. Herzog, L.E., Wang, L., Yu, E., Choi, S., Farsi, Z., Song, B.J., Pan, J.Q., and Sheng, M. (2023). Mouse mutants in schizophrenia risk genes GRIN2A and AKAP11 show EEG abnormalities in common with schizophrenia patients. Transl Psychiatry 13, 92. 10.1038/s41398-023-02393-7.

16. Camp, C.R., Vlachos, A., Klockner, C., Krey, I., Banke, T.G., Shariatzadeh, N., Ruggiero, S.M., Galer, P., Park, K.L., Caccavano, A., et al. (2023). Loss of Grin2a causes a transient delay in the electrophysiological maturation of hippocampal parvalbumin interneurons. Commun Biol 6, 952. 10.1038/s42003-023-05298-9.

17. van Os, J., and Kapur, S. (2009). Schizophrenia. Lancet 374, 635–645. 10.1016/S0140-6736(09)60995-8.

18. Abi-Dargham, A. (2014). Schizophrenia: overview and dopamine dysfunction. J Clin Psychiatry 75, e31. 10.4088/JCP.13078tx2c.

19. Grace, A.A. (2016). Dysregulation of the dopamine system in the pathophysiology of schizophrenia and depression. Nat Rev Neurosci 17, 524–532. 10.1038/nrn.2016.57.

20. Kapur, S., and Seeman, P. (2001). Does fast dissociation from the dopamine d(2) receptor explain the action of atypical antipsychotics?: A new hypothesis. Am J Psychiatry 158, 360–369. 10.1176/appi.ajp.158.3.360.

21. Matthysse, S. (1973). Antipsychotic drug actions: a clue to the neuropathology of schizophrenia? Fed Proc 32, 200–205.

22. Seeman, M.V. (2021). History of the dopamine hypothesis of antipsychotic action. World J Psychiatry 11, 355–364. 10.5498/wjp.v11.i7.355.

23. Seeman, P., Lee, T., Chau-Wong, M., and Wong, K. (1976). Antipsychotic drug doses and neuroleptic/dopamine receptors. Nature 261, 717–719. 10.1038/261717a0.

24. Abi-Dargham, A., Gil, R., Krystal, J., Baldwin, R.M., Seibyl, J.P., Bowers, M., van Dyck, C.H., Charney, D.S., Innis, R.B., and Laruelle, M. (1998). Increased striatal dopamine transmission in schizophrenia: confirmation in a second cohort. Am J Psychiatry 155, 761–767. 10.1176/ajp.155.6.761.

25. Abi-Dargham, A., Rodenhiser, J., Printz, D., Zea-Ponce, Y., Gil, R., Kegeles, L.S., Weiss, R., Cooper, T.B., Mann, J.J., Van Heertum, R.L., et al. (2000). Increased baseline occupancy of D2 receptors by dopamine in schizophrenia. Proc Natl Acad Sci U S A 97, 8104–8109. 10.1073/pnas.97.14.8104.

26. Howes, O.D., Williams, M., Ibrahim, K., Leung, G., Egerton, A., McGuire, P.K., and Turkheimer, F. (2013). Midbrain dopamine function in schizophrenia and depression: a post-mortem and positron emission tomographic imaging study. Brain 136, 3242–3251. 10.1093/brain/awt264.

27. Bagasrawala, I., Memi, F., N, V.R., and Zecevic, N. (2017). N-Methyl d-Aspartate Receptor Expression Patterns in the Human Fetal Cerebral Cortex. Cereb Cortex 27, 5041–5053. 10.1093/cercor/bhw289.

28. Bar-Shira, O., Maor, R., and Chechik, G. (2015). Gene Expression Switching of Receptor Subunits in Human Brain Development. PLoS Comput Biol 11, e1004559. 10.1371/journal.pcbi.1004559.

29. Maussion, G., Diallo, A.B., Gigek, C.O., Chen, E.S., Crapper, L., Theroux, J.F., Chen, G.G., Vasuta, C., and Ernst, C. (2015). Investigation of genes important in neurodevelopment disorders in adult human brain. Hum Genet 134, 1037–1053. 10.1007/s00439-015-1584-z.

30. Monyer, H., Burnashev, N., Laurie, D.J., Sakmann, B., and Seeburg, P.H. (1994). Developmental and regional expression in the rat brain and functional properties of four NMDA receptors. Neuron 12, 529–540. 10.1016/0896-6273(94)90210-0.

31. Wenzel, A., Scheurer, L., Kunzi, R., Fritschy, J.M., Mohler, H., and Benke, D. (1995). Distribution of NMDA receptor subunit proteins NR2A, 2B, 2C and 2D in rat brain. Neuroreport 7, 45–48.

32. Creese, I., Burt, D.R., and Snyder, S.H. (1976). Dopamine receptor binding predicts clinical and pharmacological potencies of antischizophrenic drugs. Science 192, 481–483. 10.1126/science.3854.

33. Snyder, S.H. (1976). The dopamine hypothesis of schizophrenia: focus on the dopamine receptor. Am J Psychiatry 133, 197–202. 10.1176/ajp.133.2.197.

34. Yun, S., Yang, B., Anair, J.D., Martin, M.M., Fleps, S.W., Pamukcu, A., Yeh, N.H., Contractor, A., Kennedy, A., and Parker, J.G. (2023). Antipsychotic drug efficacy correlates with the modulation of D1 rather than D2 receptor-expressing striatal projection neurons. Nat Neurosci 26, 1417–1428. 10.1038/s41593-023-01390-9.

35. Angrist, B., Rotrosen, J., and Gershon, S. (1980). Responses to apomorphine, amphetamine, and neuroleptics in schizophrenic subjects. Psychopharmacology (Berl) 67, 31–38. 10.1007/BF00427592.

36. Abi-Dargham, A., van de Giessen, E., Slifstein, M., Kegeles, L.S., and Laruelle, M. (2009). Baseline and amphetamine-stimulated dopamine activity are related in drug-naive schizophrenic subjects. Biol Psychiatry 65, 1091–1093. 10.1016/j.biopsych.2008.12.007.

37. Laruelle, M., Abi-Dargham, A., van Dyck, C.H., Gil, R., D’Souza, C.D., Erdos, J., McCance, E., Rosenblatt, W., Fingado, C., Zoghbi, S.S., et al. (1996). Single photon emission computerized tomography imaging of amphetamine-induced dopamine release in drug-free schizophrenic subjects. Proc Natl Acad Sci U S A 93, 9235–9240. 10.1073/pnas.93.17.9235.

38. Egerton, A., Chaddock, C.A., Winton-Brown, T.T., Bloomfield, M.A., Bhattacharyya, S., Allen, P., McGuire, P.K., and Howes, O.D. (2013). Presynaptic striatal dopamine dysfunction in people at ultra-high risk for psychosis: findings in a second cohort. Biol Psychiatry 74, 106–112. 10.1016/j.biopsych.2012.11.017.

39. Watanabe, M., Inoue, Y., Sakimura, K., and Mishina, M. (1992). Developmental changes in distribution of NMDA receptor channel subunit mRNAs. Neuroreport 3, 1138–1140. 10.1097/00001756-199212000-00027.

40. Kraeuter, A.K., Guest, P.C., and Sarnyai, Z. (2019). The Y-Maze for Assessment of Spatial Working and Reference Memory in Mice. Methods Mol Biol 1916, 105–111. 10.1007/978-1-4939-8994-2_10.

41. Montgomery, K.C. (1955). The relation between fear induced by novel stimulation and exploratory behavior. J Comp Physiol Psychol 48, 254–260. 10.1037/h0043788.

42. Broadhurst, P.L. (1969). Psychogenetics of emotionality in the rat. Ann N Y Acad Sci 159, 806–824. 10.1111/j.1749-6632.1969.tb12980.x.

43. Powell, C.M., and Miyakawa, T. (2006). Schizophrenia-relevant behavioral testing in rodent models: a uniquely human disorder? Biol Psychiatry 59, 1198–1207. 10.1016/j.biopsych.2006.05.008.

44. Bondi, C., Matthews, M., and Moghaddam, B. (2012). Glutamatergic animal models of schizophrenia. Curr Pharm Des 18, 1593–1604. 10.2174/138161212799958576.

45. McCarthy, J.M., Treadway, M.T., Bennett, M.E., and Blanchard, J.J. (2016). Inefficient effort allocation and negative symptoms in individuals with schizophrenia. Schizophr Res 170, 278–284. 10.1016/j.schres.2015.12.017.

46. Treadway, M.T., Peterman, J.S., Zald, D.H., and Park, S. (2015). Impaired effort allocation in patients with schizophrenia. Schizophr Res 161, 382–385. 10.1016/j.schres.2014.11.024.

47. Kirschner, M., Hager, O.M., Bischof, M., Hartmann-Riemer, M.N., Kluge, A., Seifritz, E., Tobler, P.N., and Kaiser, S. (2016). Deficits in context-dependent adaptive coding of reward in schizophrenia. NPJ Schizophr 2, 16020. 10.1038/npjschz.2016.20.

48. Bradshaw, C.M., and Killeen, P.R. (2012). A theory of behaviour on progressive ratio schedules, with applications in behavioural pharmacology. Psychopharmacology (Berl) 222, 549–564. 10.1007/s00213-012-2771-4.

49. Reddy, L.F., Waltz, J.A., Green, M.F., Wynn, J.K., and Horan, W.P. (2016). Probabilistic Reversal Learning in Schizophrenia: Stability of Deficits and Potential Causal Mechanisms. Schizophr Bull 42, 942–951. 10.1093/schbul/sbv226.

50. Culbreth, A.J., Gold, J.M., Cools, R., and Barch, D.M. (2016). Impaired Activation in Cognitive Control Regions Predicts Reversal Learning in Schizophrenia. Schizophr Bull 42, 484–493. 10.1093/schbul/sbv075.

51. Kim, Y., Wood, J., and Moghaddam, B. (2012). Coordinated activity of ventral tegmental neurons adapts to appetitive and aversive learning. PLoS One 7, e29766. 10.1371/journal.pone.0029766.

52. Kim, Y.B., Matthews, M., and Moghaddam, B. (2010). Putative gamma-aminobutyric acid neurons in the ventral tegmental area have a similar pattern of plasticity as dopamine neurons during appetitive and aversive learning. Eur J Neurosci 32, 1564–1572. 10.1111/j.1460-9568.2010.07371.x.

53. Lefner, M.J., and Moghaddam, B. (2025). Flexible updating of reward and punishment contingencies by VTA GABA neurons. Curr Biol. 10.1016/j.cub.2025.07.021.

54. Lubow, R.E. (2005). Construct validity of the animal latent inhibition model of selective attention deficits in schizophrenia. Schizophr Bull 31, 139–153. 10.1093/schbul/sbi005.

55. Lubow, R.E., and Gewirtz, J.C. (1995). Latent inhibition in humans: data, theory, and implications for schizophrenia. Psychol Bull 117, 87–103. 10.1037/0033-2909.117.1.87.

56. Laruelle, M., and Abi-Dargham, A. (1999). Dopamine as the wind of the psychotic fire: new evidence from brain imaging studies. J Psychopharmacol 13, 358–371. 10.1177/026988119901300405.

57. Howes, O.D., Kambeitz, J., Kim, E., Stahl, D., Slifstein, M., Abi-Dargham, A., and Kapur, S. (2012). The nature of dopamine dysfunction in schizophrenia and what this means for treatment. Arch Gen Psychiatry 69, 776–786. 10.1001/archgenpsychiatry.2012.169.

58. Millard, S.J., Bearden, C.E., Karlsgodt, K.H., and Sharpe, M.J. (2022). The prediction- error hypothesis of schizophrenia: new data point to circuit-specific changes in dopamine activity. Neuropsychopharmacology 47, 628–640. 10.1038/s41386-021-01188-y.

59. Katthagen, T., Kaminski, J., Heinz, A., Buchert, R., and Schlagenhauf, F. (2020). Striatal Dopamine and Reward Prediction Error Signaling in Unmedicated Schizophrenia Patients. Schizophr Bull 46, 1535–1546. 10.1093/schbul/sbaa055.

60. Sun, F., Zeng, J., Jing, M., Zhou, J., Feng, J., Owen, S.F., Luo, Y., Li, F., Wang, H., Yamaguchi, T., et al. (2018). A Genetically Encoded Fluorescent Sensor Enables Rapid and Specific Detection of Dopamine in Flies, Fish, and Mice. Cell 174, 481–496 e419. 10.1016/j.cell.2018.06.042.

61. Bromberg-Martin, E.S., Matsumoto, M., and Hikosaka, O. (2010). Dopamine in motivational control: rewarding, aversive, and alerting. Neuron 68, 815–834. 10.1016/j.neuron.2010.11.022.

62. Ohi, K., Shimada, T., Nitta, Y., Kihara, H., Okubo, H., Uehara, T., and Kawasaki, Y. (2016). Specific gene expression patterns of 108 schizophrenia-associated loci in cortex. Schizophr Res 174, 35–38. 10.1016/j.schres.2016.03.032.

63. Howes, O.D., Montgomery, A.J., Asselin, M.C., Murray, R.M., Valli, I., Tabraham, P., Bramon-Bosch, E., Valmaggia, L., Johns, L., Broome, M., et al. (2009). Elevated striatal dopamine function linked to prodromal signs of schizophrenia. Arch Gen Psychiatry 66, 13–20. 10.1001/archgenpsychiatry.2008.514.

64. Howes, O.D., and Nour, M.M. (2016). Dopamine and the aberrant salience hypothesis of schizophrenia. World Psychiatry 15, 3–4. 10.1002/wps.20276.

65. Fletcher, P.C., and Frith, C.D. (2009). Perceiving is believing: a Bayesian approach to explaining the positive symptoms of schizophrenia. Nat Rev Neurosci 10, 48–58. 10.1038/nrn2536.

66. Olney, J.W., and Farber, N.B. (1995). Glutamate receptor dysfunction and schizophrenia. Arch Gen Psychiatry 52, 998–1007. 10.1001/archpsyc.1995.03950240016004.

67. Morris, R.W., Vercammen, A., Lenroot, R., Moore, L., Langton, J.M., Short, B., Kulkarni, J., Curtis, J., O’Donnell, M., Weickert, C.S., and Weickert, T.W. (2012). Disambiguating ventral striatum fMRI-related BOLD signal during reward prediction in schizophrenia. Mol Psychiatry 17, 235, 280–239. 10.1038/mp.2011.75.

68. Corlett, P.R., Murray, G.K., Honey, G.D., Aitken, M.R., Shanks, D.R., Robbins, T.W., Bullmore, E.T., Dickinson, A., and Fletcher, P.C. (2007). Disrupted prediction-error signal in psychosis: evidence for an associative account of delusions. Brain 130, 2387–2400. 10.1093/brain/awm173.

69. Schultz, W., Dayan, P., and Montague, P.R. (1997). A neural substrate of prediction and reward. Science 275, 1593–1599. 10.1126/science.275.5306.1593.

70. Schultz, W. (1998). Predictive reward signal of dopamine neurons. J Neurophysiol 80, 1–27. 10.1152/jn.1998.80.1.1.

71. Balu, D.T., and Coyle, J.T. (2015). The NMDA receptor ’glycine modulatory site’ in schizophrenia: D-serine, glycine, and beyond. Curr Opin Pharmacol 20, 109–115. 10.1016/j.coph.2014.12.004.

72. Moghaddam, B., and Adams, B.W. (1998). Reversal of phencyclidine effects by a group II metabotropic glutamate receptor agonist in rats. Science 281, 1349–1352. 10.1126/science.281.5381.1349.

73. Kinon, B.J., Millen, B.A., Zhang, L., and McKinzie, D.L. (2015). Exploratory analysis for a targeted patient population responsive to the metabotropic glutamate 2/3 receptor agonist pomaglumetad methionil in schizophrenia. Biol Psychiatry 78, 754–762. 10.1016/j.biopsych.2015.03.016.

74. Wager-Miller, J., Murphy Green, M., Shafique, H., and Mackie, K. (2020). Collection of Frozen Rodent Brain Regions for Downstream Analyses. J Vis Exp. 10.3791/60474.

75. Paxinos, G., & Watson, C. (1998). A stereotaxic atlas of the rat brain. (New York: Academic).

76. Luck, C., Haitjema, C., and Heger, C. (2021). Simple Western: Bringing the Western Blot into the Twenty-First Century. Methods Mol Biol 2261, 481–488. 10.1007/978-1-0716-1186-9_30.

77. Hunker, A.C., Soden, M.E., Krayushkina, D., Heymann, G., Awatramani, R., and Zweifel, L.S. (2020). Conditional Single Vector CRISPR/SaCas9 Viruses for Efficient Mutagenesis in the Adult Mouse Nervous System. Cell Rep 30, 4303–4316 e4306. 10.1016/j.celrep.2020.02.092.

78. Richardson, N.R., and Roberts, D.C. (1996). Progressive ratio schedules in drug self-administration studies in rats: a method to evaluate reinforcing efficacy. J Neurosci Methods 66, 1–11. 10.1016/0165-0270(95)00153-0.

79. Rescorla, R.A.W., A.R. (1972). A theory of Pavlovian conditioning: Variations in the effectiveness of reinforcement and nonreinforcement. In Classical conditioning II: Current research and theory, A.H.P. Black, W. F., ed. (Appleton-Century-Crofts), pp. 64–99.

80. Pearce, J.M., and Hall, G. (1980). A model for Pavlovian learning: variations in the effectiveness of conditioned but not of unconditioned stimuli. Psychol Rev 87, 532–552.

81. Roesch, M.R., Esber, G.R., Li, J., Daw, N.D., and Schoenbaum, G. (2012). Surprise! Neural correlates of Pearce-Hall and Rescorla-Wagner coexist within the brain. Eur J Neurosci 35, 1190–1200. 10.1111/j.1460-9568.2011.07986.x.

