## supplement figures for "Reduction of *Grin2a* in adolescent rat dopamine neurons confers a phenotype relevant to psychosis"

**Supplemental Figures**

a

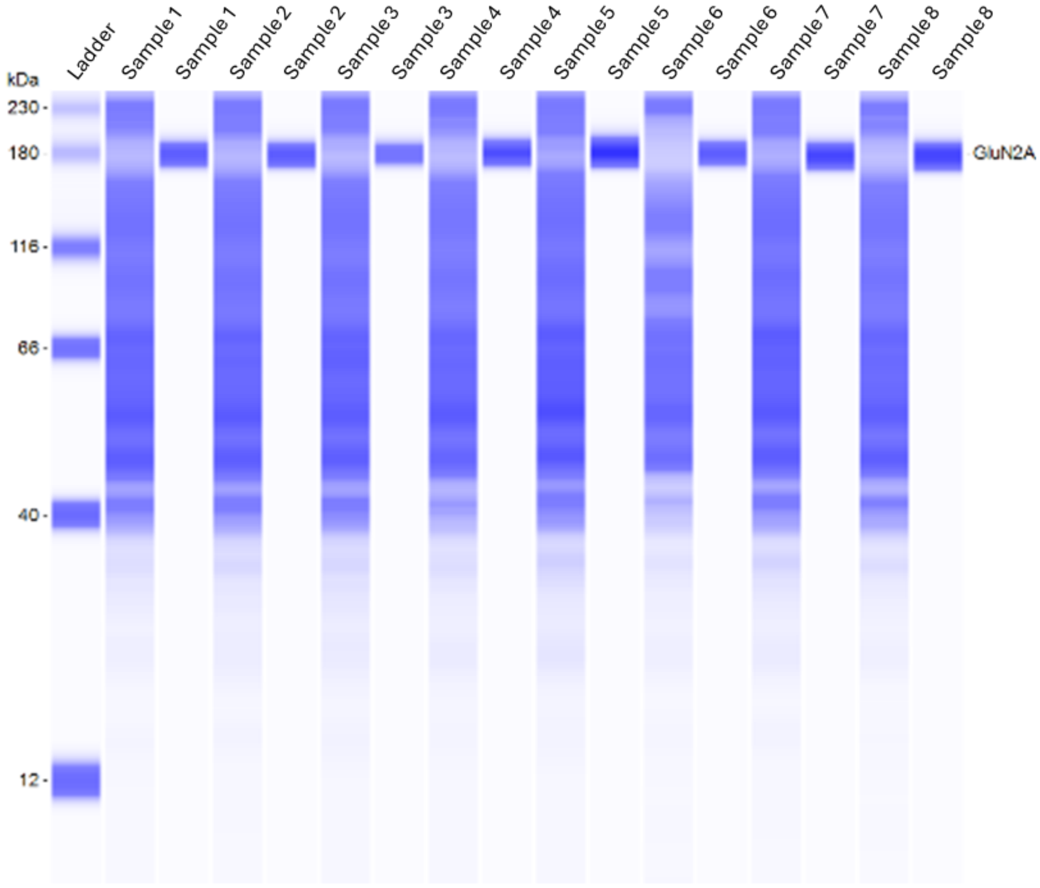


**Figure S1: Representative pseudo-blot from Wes**

**a,** Example pseudo-blot for protein analysis of neural tissue punches with automated Western blotting experiments from the hippocampus. Each sample was run in two columns, one to detect total protein for normalization, the other to stain for GluN2A (~170kDa).

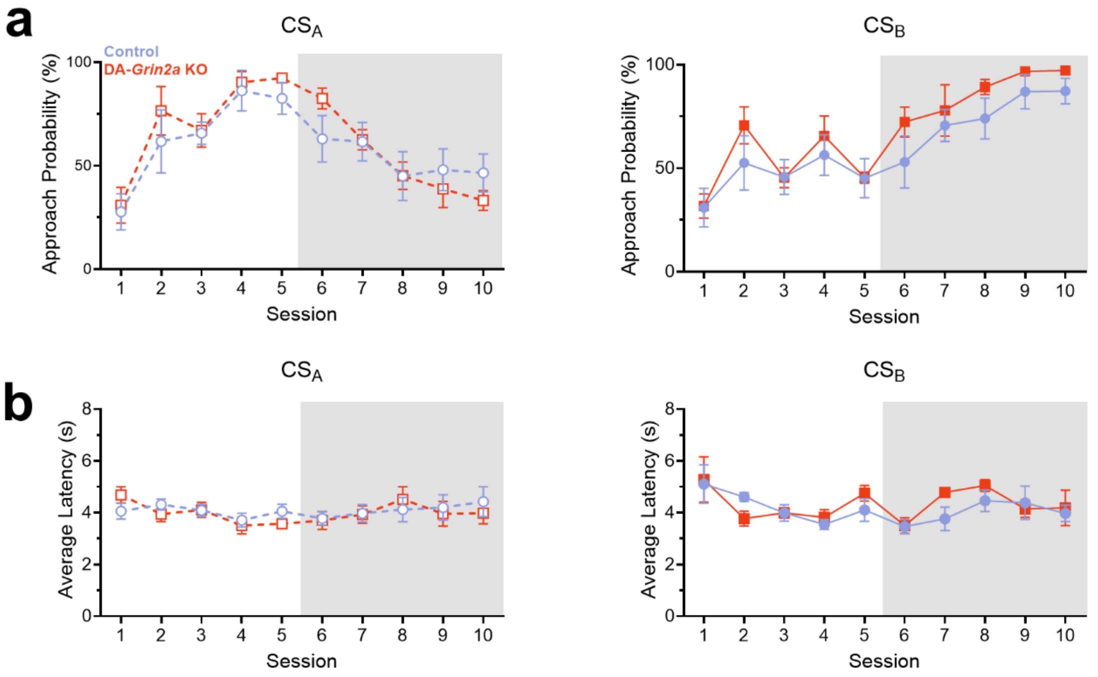


**Figure S2: Additional analysis of adolescent behavior during FCL**

**a,** Approach probability to the food port during $CS_A$ (left) or $CS_B$ (right) presentation. **b,** Average latency to approach the food port during $CS_A$ (left) or $CS_B$ (right) presentation. Error bars denote ±SEM.

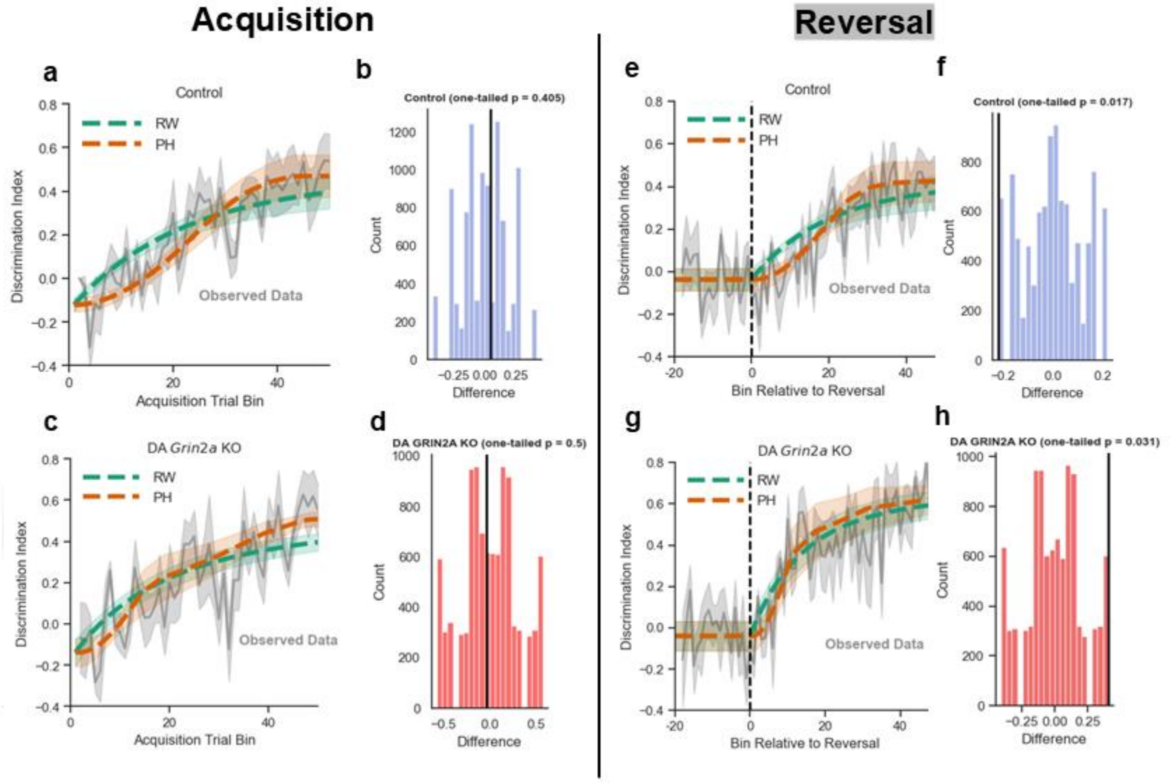


**Figure S3: Simulation results and permutation tests for Pearce-Hall (PH) and Rescorla-Wagner (RW) learning models for adolescent FCL behavior**

**a,** Binned discrimination index data (grey, 5 trial bins) over acquisition and group averaged predictions of PH (orange) and RW (green) models for the CS associated with reward. Both models captured conditioned approach behavior over acquisition training in Control animals. **b,** Null distribution from permutation tests based on difference in fit between PH and RW models. No model was clearly preferred (*p*=0.40). Solid black line indicates observed difference in fit between the models (positive = RW preferred, negative = PH preferred). **c,** Same as **a**, but for DA-*Grin2a* KO acquisition data. **d,** Permutation test null distribution and observed difference (black) for DA-*Grin2a* KO acquisition data. Like controls, neither model was clearly preferred (*p*=0.50)**. e,** Control group binned discrimination index data (grey, 5 trial bins) for two sessions before and all sessions after contingency reversal for the CS which switched from punishment to reward predictive. Group averaged predictions from PH (orange) and RW (green) are indicated by dashed lines. The RW model generally overestimated behavior in early reversal. **f,** Null distribution from permutation tests based on difference in fit between PH and RW models. Solid black line indicates observed difference in fit between the models (positive = RW preferred, negative = PH preferred). PH learning mechanisms were generally favored compared to RW (*p*=0.017). **g,** Same as **e,** but for DA-*Grin2a* KO animals. Unlike controls, behavior at reversal was well accounted by RW models. **h,** same as **f,** but for DA-*Grin2a* KO animals. RW learning mechanisms were generally favored over PH for the DA-*Grin2a* KO group (*p*=0.03). Line shading denotes ± SEM.

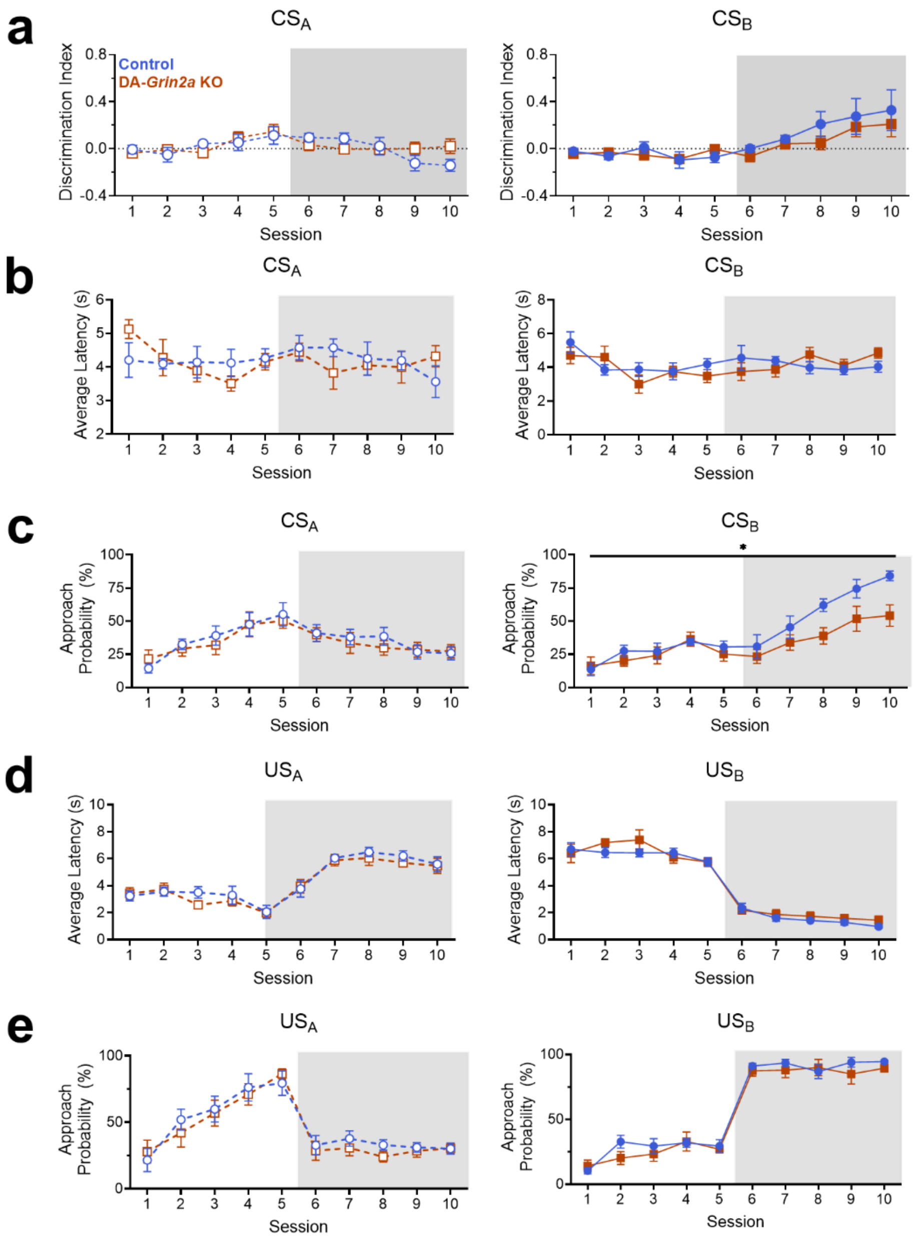


**Figure S4: Analysis of tethered young adult behavior during FCL**

**a,** Discrimination index for $CS_A$ (left) or $CS_B$ (right) **b,** Latency to approach the food port following $CS_A$ (left) or $CS_B$ (right) onset. **c,** Probability to approach the food port during CS presentation. Control animals were significantly more likely to approach the food port in response to $CS_B$ compared to DA-*Grin2a* KO animals (*$p = 0.0389$). **d,** Latency to approach the food port following US onset. **e,** Probability to approach the food port following US onset. Error bars denote ±SEM.

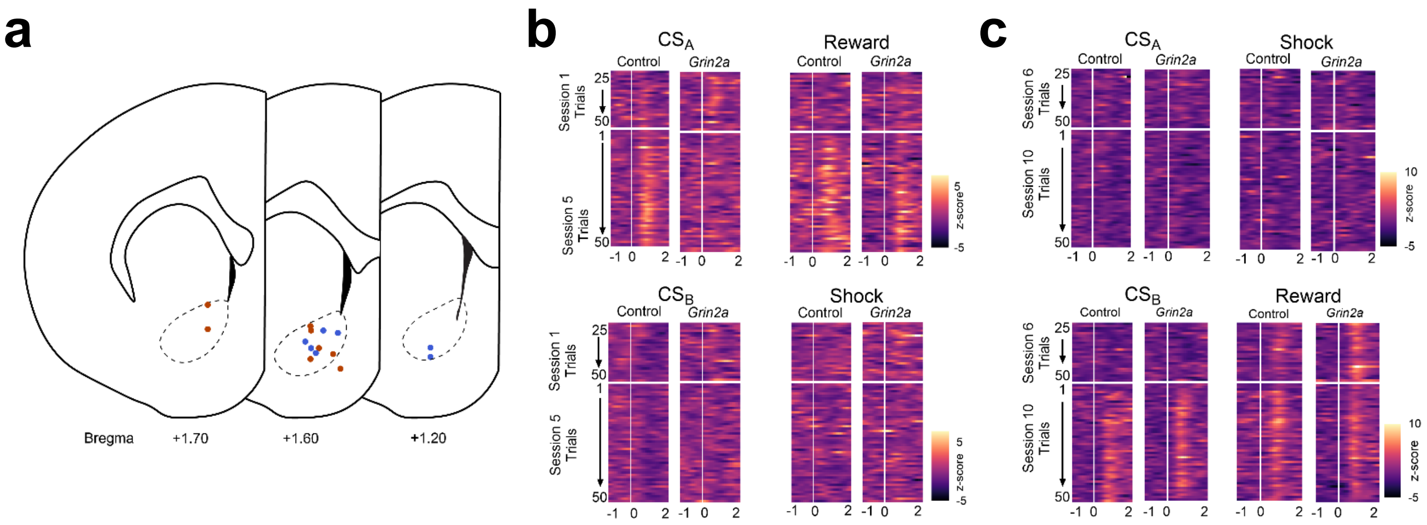


**Figure S5: Fiber photometry hit map and representative heatmaps**

**a,** Location of fiber placement for fiber photometry experiments. Orange dots reflect DA-*Grin2a* KO animals, blue dots reflect control animals. **b,** Representative heatmaps of single control and single DA-*Grin2a* KO rats' dopaminergic response to CS onset and reward/shock delivery during initial association learning. **c,** Representative heatmaps of control and DA-*Grin2a* KO rats' dopaminergic response to CS onset and reward/shock delivery following the contingency switch.

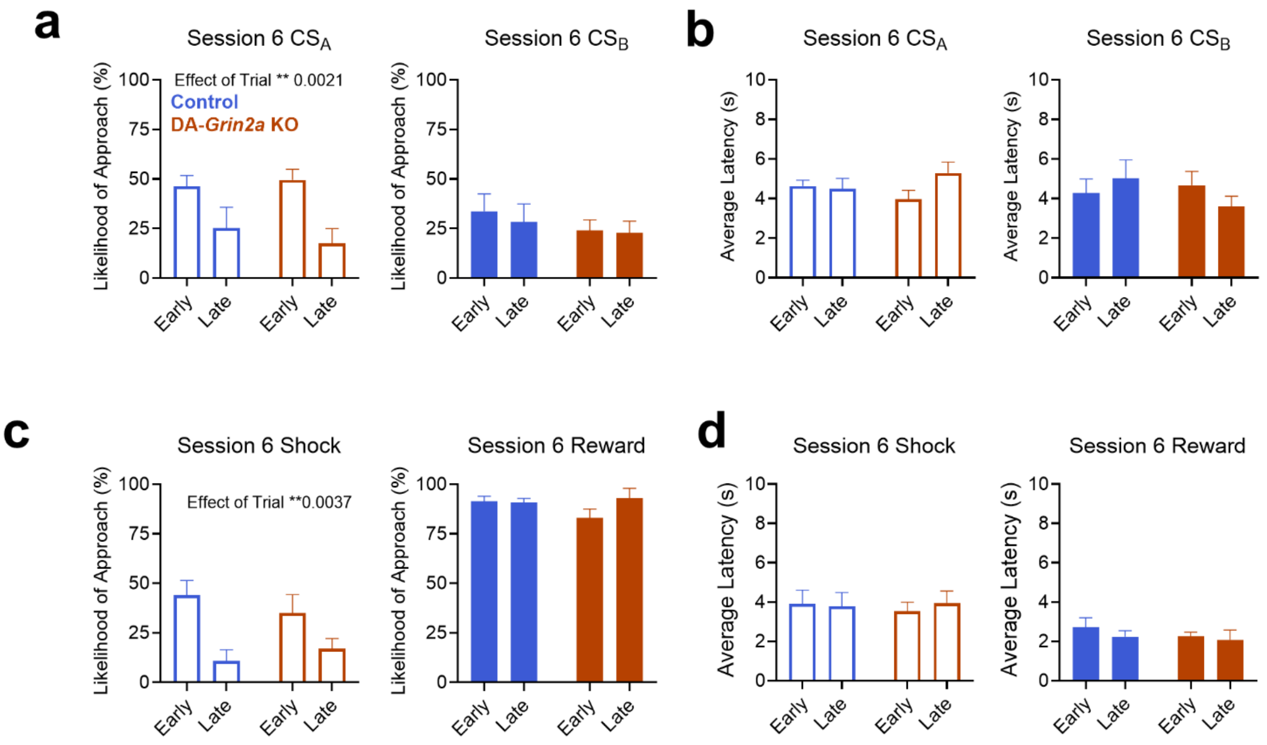


**Figure S6: Behavioral analysis of Early and Late trials of FCL Session 6 in young adult animals.**

**a,** Probability to approach food port following $CS_A$ (left) or $CS_B$ (right) onset. **b**, Average latency to approach food port following $CS_A$ (left) or $CS_B$ (right) onset**. c,** Likelihood to approach the food port following US onset. **d,** Average latency to approach food port following US onset. Error bars denote +SEM.

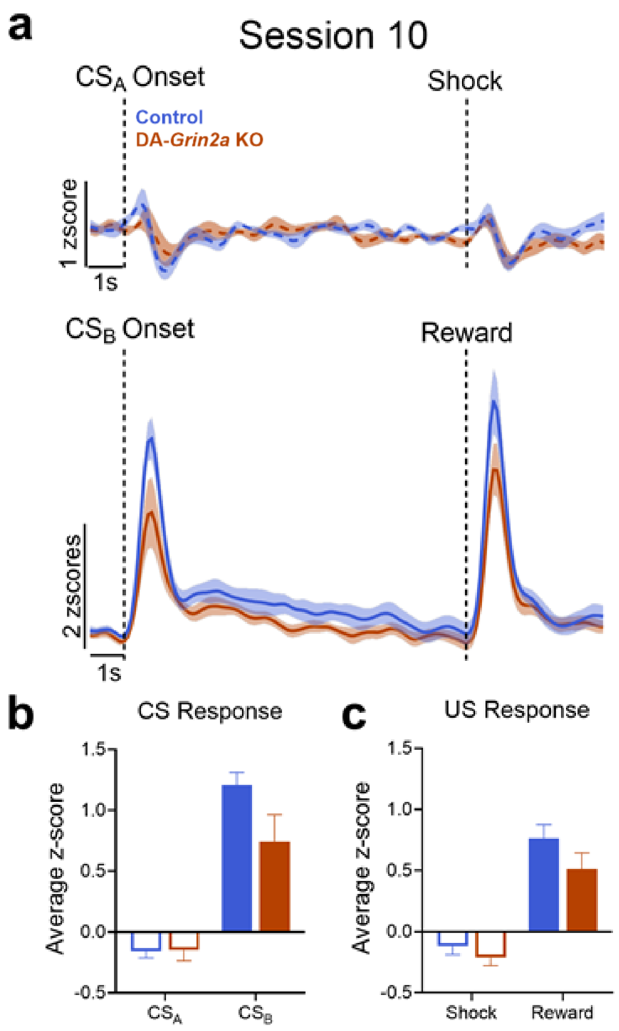


**Figure S7: Fiber photometry analysis of Session 10 of FCL**

**a,** Fluorescent signal in $CS_A$ and $CS_B$ trials on Session 10 of FCL. **b,** Quantification of average z-score across 2 s period at CS delivery showed negative responses to the punishment-predicting cue ($CS_A$) and positive responses to the reward-predicting cue ($CS_B$) in both groups. **c,** Quantification of average z-score across 4 s period after US onset showed a similar pattern. Error bars and shading denote ±SEM.
